# Compensatory ion transport buffers daily protein rhythms to regulate osmotic balance and cellular physiology

**DOI:** 10.1101/2020.05.28.118398

**Authors:** Alessandra Stangherlin, David C. S. Wong, Silvia Barbiero, Joseph L. Watson, Aiwei Zeng, Estere Seinkmane, Sew Peak Chew, Andrew D. Beale, Edward A. Hayter, Alina Guna, Alison J. Inglis, Eline Bartolami, Stefan Matile, Nicolas Lequeux, Thomas Pons, Jason Day, Gerben van Ooijen, Rebecca M. Voorhees, David A. Bechtold, Emmanuel Derivery, Rachel S. Edgar, Peter Newham, John S. O’Neill

## Abstract

Between 6-20% of the cellular proteome is under circadian control to tune cell function with cycles of environmental change. For cell viability, and to maintain volume within narrow limits, the osmotic pressure exerted by changes in the soluble proteome must be compensated. The mechanisms and consequences underlying compensation are not known. Here, we show in cultured mammalian cells and *in vivo* that compensation requires electroneutral active transport of Na^+^, K^+^, and Cl^−^ through differential activity of SLC12A family cotransporters. In cardiomyocytes *ex vivo* and *in vivo*, compensatory ion fluxes alter their electrical activity at different times of the day. Perturbation of soluble protein abundance has commensurate effects on ion composition and cellular function across the circadian cycle. Thus, circadian regulation of the proteome impacts ion homeostasis with substantial consequences for the physiology of electrically active cells such as cardiomyocytes.

## Introduction

The abundance of many proteins exhibits ~24 h rhythms, regulated by cell autonomous circadian timing mechanisms that align physiology with the day-night cycle^1–6^. Between 6% and 20% of cellular proteins are under circadian control, and the expression of most oscillating proteins peaks during translational “rush hours” that typically coincide with the organism’s habitual active phase^1–7^. Daily regulation of mechanistic target-of-rapamycin complexes (mTORC) partitions phases of increased protein production (increased mTORC activity) from those of increased catabolism (decreased mTORC activity)^8–11^. This temporal organisation of protein homeostasis (proteostasis) permits the most efficient use of bioenergetic resources, both *in vivo* and in cultured mammalian cells^8–10,12–14^. How cellular protein concentration varies over circadian time in different intracellular compartments is currently unknown.

In the crowded cytosol, macromolecules (300-550 mg/mL^15^) and K^+^ ions (~145 mM^16^) are the major determinants of cytosolic osmotic potential, balanced by high extracellular concentrations of Na^+^ and Cl^−^ ions. Cells must therefore accommodate any daily variation in cytosolic protein abundance without compromising osmotic homeostasis (osmostasis), which would have deleterious effects on cellular function and viability^17–20^. Several osmoregulatory mechanisms are in place to protect cells from changes in extracellular tonicity^17,21^, but very little is known about how cells compensate for physiological changes in intracellular macromolecule content to remain in osmotic equilibrium during the circadian cycle.

Electroneutral cotransporters of the SLC12A family are ubiquitously-expressed symporters that regulate transmembrane ion gradients and cell volume across a wide variety of tissues^22^. They couple ion transport with the respective transmembrane Na^+^ and K^+^ gradients, that are ultimately established by the ubiquitous and essential Na/K-ATPase antiporter^17^. Na-K-Cl (NKCC) and NaCl (NCC) cotransporters facilitate electroneutral secondary active ion influx (1 Na^+^, 1 K^+^, with 2 Cl^−^ and 1 Na^+^ with 1 Cl^−^, respectively), whereas K-Cl (KCC) transporters facilitate electroneutral ion efflux (1 K^+^ and 1 Cl^−^)^22^. Changes in the relative activity of NKCC and KCC isoforms regulate intracellular ion levels, in particular Cl^−^. During neuronal development, for example, progressive expression of KCC2 increases intracellular Cl^−^ and mediates the transition between excitatory and inhibitory GABA signalling^23,24^. Furthermore, differential KCC/NKCC activity is thought to mediate GABAergic synaptic plasticity of circadian regulation in the mammalian suprachiasmatic nucleus^25,26^.

SLC12A cotransporters show high amplitude circadian rhythms at the mRNA level^4,27^, but their activity is primarily regulated post-translationally via a coordinated network of kinases, including the essential serine/threonine kinase oxidative stress response kinase 1 (OXSR1)^17,28^. A large body of evidence supports the differential regulation of SLC12A cotransporters by OXSR1: phosphorylation of NKCC and NCC activates ion import (Na^+^, K^+^, Cl^−^) whilst phosphorylation of KCC attenuates ion export (K^+^ and Cl^−^)^28,29^. The activity of OXSR1 itself is regulated upstream through phosphorylation by lysine deficient protein kinases family members (WNKs), which are stimulated by molecular crowding and low cytosolic Cl^−28–30^. Daily variation in the phosphorylation of WNK isoforms has been reported in mouse liver ^12^ and forebrain^27^, as has the phosphorylation of several members of the SLC12A family^27,31^.

Here, we describe a fundamental and isovolumetric process whereby electroneutral ion transport buffers intracellular osmotic potential against daily and acute variations in protein abundance. Consequently, the ionic composition of the cell is not constant throughout the day, as has been assumed^32^. We found that dynamic changes in ion abundance drive oscillations in cellular physiology that impart temporal regulation to cardiomyocyte cell function and heart rate. More broadly, our data suggest that the cellular capacity for dynamic ion transport is important for protein homeostasis.

## Results

### Cell-autonomous rhythms in soluble protein abundance

Up to a fifth of the cellular proteome exhibits 24 h cycles in abundance^1–7^, due in part to circadian oscillations of mTORC1 activity, ribosome biogenesis, and global protein synthesis rates^10,33,34^. We confirmed that mTORC1 activity rhythms occur cell-autonomously in quiescent adult mouse fibroblasts, maintained under constant conditions (37°C), following synchronisation by daily temperature cycles (12 h:12 h 32°C:37°C) (Fig. 1a, Supplementary Fig. 1a).

**Fig. 1.**
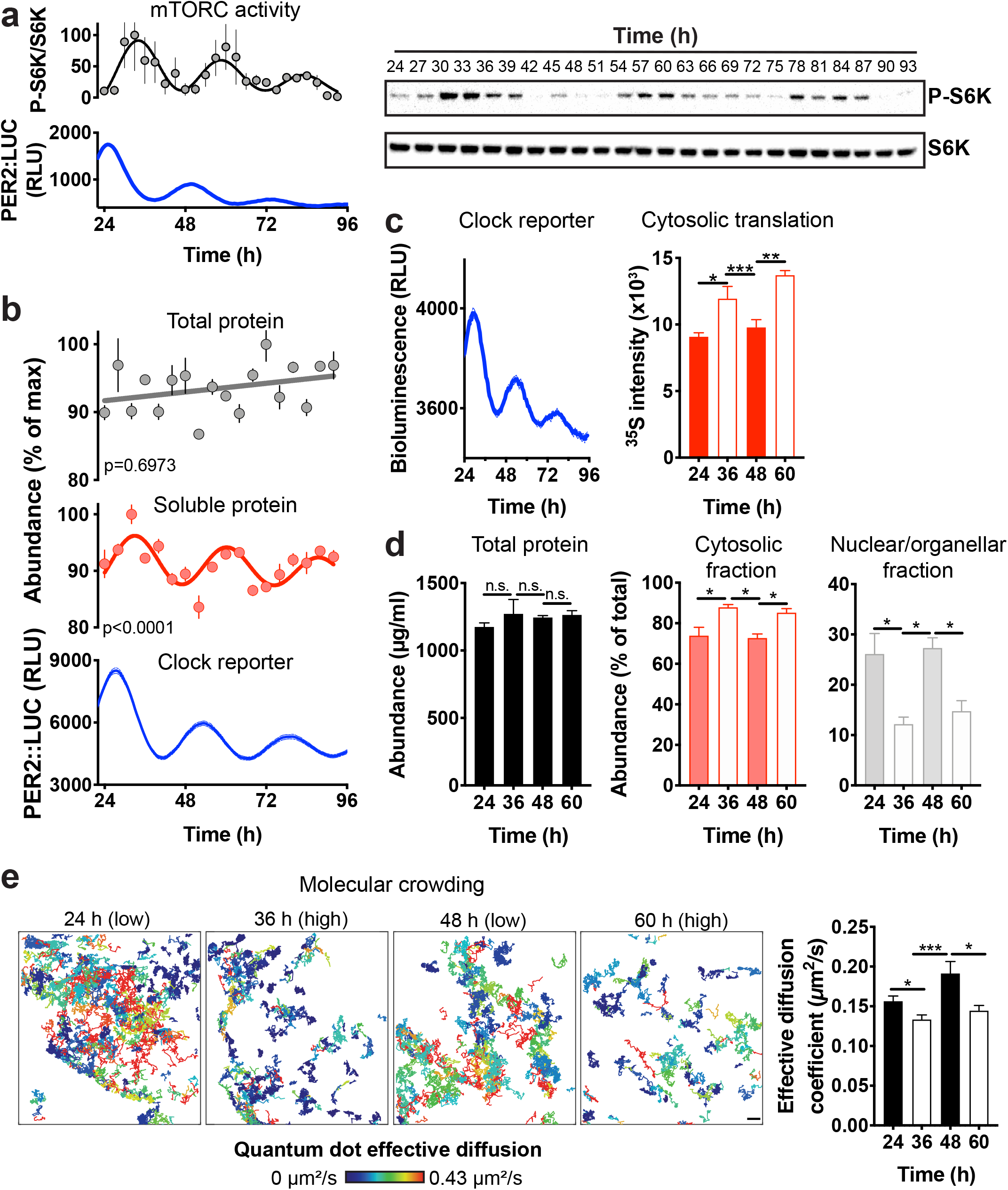
Circadian variation in cytosolic protein content in mouse fibroblasts. (**a**) Cell-autonomous circadian mTORC1 activity detected by immunoblots of phospho-S6 kinase and S6 kinase abundance in fibroblasts sampled every 3 h for three days in constant conditions (n=3) (right). Blot quantification and parallel PER2::LUC bioluminescent clock reporter (left). (**b**) Total and soluble protein quantification of cell lysates sampled ever 4 h and parallel PER2::LUC bioluminescent clock reporter (n=3). Protein abundance values were normalised for the maximal value in each timeseries. (**c**) ^35^S-met/cys incorporation assay at peak (36 h/60 h) and trough (24 h/48 h) of S6K phosphorylation rhythms (n=4) and parallel PER2::LUC activity (n=3). (**d**) Protein quantification of total cell lysates, nuclear/organellar and cytosolic fractions at peak (36 h/60 h) and trough (24 h/48 h) of protein rhythms. (**e**) Effective diffusion rate of QDs in fibroblasts (n=81, 83, 51, 63, respectively) and representative tracking images. Colour key represents diffusion rate. Mean ± SEM shown throughout. p-values indicate comparison of damped cosine wave with straight-line fit (null hypothesis = no rhythm). Significance was calculated using one-way ANOVA and Tukey’s multi comparisons test (MCT) (**c**), one-way ANOVA and Sidak’s MCT (**d**), and Kruskal-Wallis with Dunn’s MCT (**e**).

mTORC1 activation stimulates translation^35^, cytoplasmic targeting of many different macromolecules, and molecular crowding^36^. We therefore asked whether we could detect circadian oscillations in the level of soluble cytosolic protein, since many rhythmically abundant proteins localise to this compartment. Total protein lysates and cytosolic soluble extracts (hereafter ‘soluble protein’) were prepared from fibroblast cultures over three consecutive days, under constant conditions as above (Supplementary Fig. 1a), and selective lysis of the plasma membrane by digitonin was confirmed by mass spectrometry (Supplementary Fig. 1b)^37^. While total cellular protein showed no oscillation, we detected clear ~24 h rhythms in the concentration of soluble protein extracted from cells (Fig. 1b). Peaks were observed 4-6 h after the peak of the clock reporter PER2::LUC (PERIOD 2-firefly LUCIFERASE fusion reporter^38^), coinciding with both maximal mTORC1 activity (Fig.1a) and increased translation rate in the soluble extract (Fig. 1c).

These findings were validated using cellular fractionation by differential centrifugation, following lysis by NP-40 at biological times corresponding to the peak and trough of the soluble protein rhythm (Fig. 1d). While total protein levels did not change, we observed reciprocal time-of-day variation in protein abundance between the cytosolic and nuclear/organellar fractions, compatible with a cell-autonomous daily rhythm in the sequestration of cytosolic proteins to other cellular compartments.

Diffusion of macromolecules in aqueous solution is sensitive to the concentration of other colloidal solutes, i.e. the level of macromolecular crowding is inversely proportional to their diffusion rate^39^. Therefore, to further validate the observed rhythm in cytosolic protein concentration, we measured the diffusion of inert quantum dot nanoparticles (QDs, 20 nm diameter) in the cytosol^40^. We reasoned that changes in abundance of macromolecules during the circadian cycle should inversely impact the diffusion of QDs (Supplementary Fig. 1c). In line with prediction, we observed time-of-day variation in the effective diffusion coefficient of QDs, with faster diffusion occurring at the trough of soluble protein rhythms (Fig. 1e, Supplementary Movie 1 and 2).

Importantly, mTORC inhibition with a low, sub-saturating concentration of the inhibitor torin1^41^ attenuated the peak of soluble protein rhythms (Supplementary Fig. 2a), and commensurately diminished molecular crowding reported by QD diffusion (Supplementary Fig. 2b). As would be expected, the effect of torin1 on soluble protein and crowding was most apparent at biological times when mTORC1 is maximally active, compared with 12 h later. Overall, these observations suggest that rhythms in cytosolic protein abundance and molecular crowding are at least partially dependent on cell-autonomous circadian regulation of mTORC1 activity, through some combination of rhythmic protein production and differential association of cytosolic proteins with other cellular compartments.

### Antiphasic oscillations in soluble protein and intracellular ion abundance

Most cells are highly permeable to water, and therefore susceptible to changes in volume upon osmotic challenge. Body fluid homeostasis ensures that cells are rarely subject to acute perturbations in extracellular osmolarity, but they must defend cell volume against fluctuations in intracellular osmolarity^6,17^. Without compensatory mechanisms, the high amplitude rhythm in soluble protein we observe would result in potentially deleterious oscillations in cell volume, as water moves to restore osmotic equilibrium^21^. As expected therefore, we found no commensurate rhythm in cell volume (Fig. 2a, Supplementary Fig. 3a); meaning cells successfully buffer osmotic potential against gradual changes in cytosolic protein concentration over circadian timeframes.

**Fig. 2.**
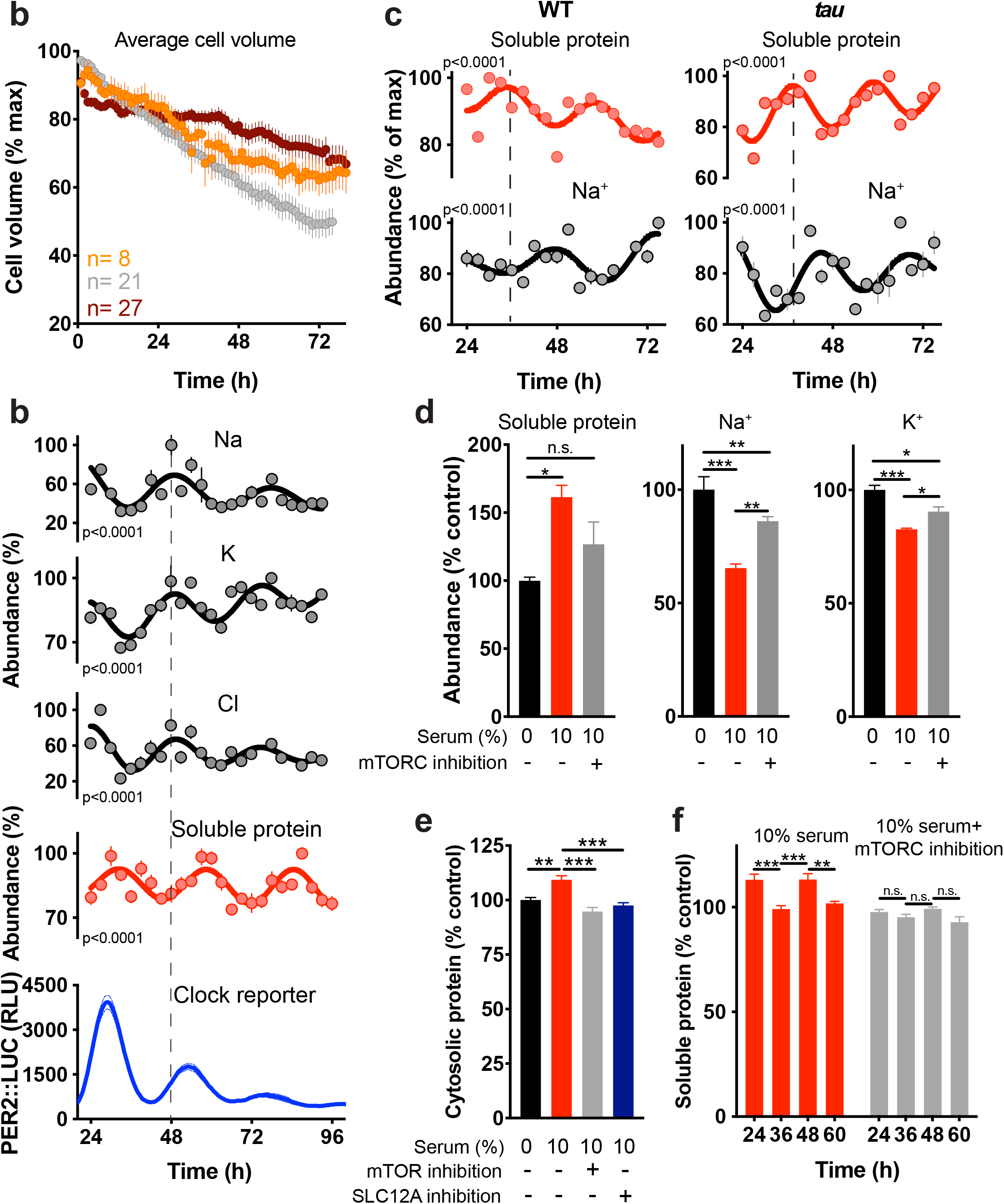
Ion flux buffers changes in cytosolic protein abundance in mouse fibroblasts. (**a**) Mean cell volume of fibroblasts across time from three independent recordings. Average cell volume data were normalised for the largest value within each replicate. (**b**) Abundance of selected ions (n=4), soluble protein (n=3), and PER2::LUC bioluminescence recordings (n=3). Abundance values were normalised for the maximal value in each timeseries. (**c**) Quantification of intracellular Na^+^ (n=4) and labile cytosolic protein abundance (n=3) in WT and *tau* mutant fibroblasts under constant conditions over two days. (**d**) % change in soluble protein abundance (n=3) and Na^+^ and K^+^ abundance (n=4) in unsynchronised cells upon 6 h treatment with 10% serum ± 50 nM mTOR inhibitor torin1. (**e**) % change in soluble protein upon treatment with 10% serum ± 50 nM mTOR inhibitor torin1 or 10% serum ± 50 μM DIOA and 100 μM Bumetanide (n=3) (**f**) % fold increase in soluble protein upon 4 h treatment with 10% serum ± 50 nM mTOR inhibitor torin1at indicated times (n=4). Mean ± SEM shown throughout. p-values in (**b**) and (**c**) indicate comparison of damped cosine wave with straight-line fit (null hypothesis = no rhythm). Statistical tests are one-way ANOVA and Dunnet’s MCT (**d)** one-away ANOVA and Tukey **(e**) and two-way ANOVA and Sidak’s MCT (**f**).

In light of previous observations^9,42^, we hypothesised that transmembrane ion flux was responsible for this isovolumetric compensatory mechanism. We focused on intracellular metal ions, the most abundant osmotically active species in cells, and analysed the intracellular ionic composition of primary fibroblasts across three circadian cycles by inductively-coupled plasma mass spectrometry (ICP-MS). Importantly, we observed ~24 h cellular ion rhythms for intracellular K^+^, Na^+^ and Cl^−^, which oscillated in antiphase with soluble protein (Fig. 2b). Several other biologically-relevant ions were not similarly rhythmic (Supplementary Fig. 3b).

To assess whether the ion and cytosolic protein rhythms are under cell-autonomous circadian control, we analysed cells from short period *tau* mutant mice^43^, which express circadian rhythms that run faster than wild type. Soluble protein and ion abundance oscillated in antiphase, as with wild type cells, but showed correspondingly shorter period oscillations (~21 h) (Fig. 2c, Supplementary Fig. 3c). In a parallel investigation, on a different clock mutant cell line, we found that increased amplitude of soluble protein levels resulted in a decreased relative amplitude of K^+^ oscillations as compared to WT cells^44^. These data suggest that cell-autonomous circadian clock mechanisms coordinate antiphasic oscillations of cytosolic ion and protein content.

### Reciprocal regulation of soluble protein and intracellular ion abundance

Our data suggest that reciprocal changes in ion transport compensate for changes in soluble protein concentration in order to maintain cellular osmostasis. We tested this relationship by treating unsynchronised cells with a bolus of serum to stimulate mTORC activity and thereby increase soluble protein levels in the cytosol^45^. We found acute addition of serum induced an mTORC-dependent increase in soluble protein with a commensurate decrease in ion levels (Fig. 2d). It follows that the ability of a cell to accommodate increases in cytosolic protein may be limited by its capacity to effect compensatory ion export. To test this, we acutely stimulated cells with serum in the presence of inhibitors of SLC12A family members, NKCC and KCC, cotransporters that control the movement of Na^+^, K^+^, and Cl^−22,46^. Importantly, we found that the inhibition of ion transport abolished the serum-induced protein increase as effectively as the mTORC inhibition (Fig. 2e).

Because intracellular ion abundance varies across the day, we predicted that at biological times when ion abundance is low, the reduced cellular capacity to buffer osmotic potential should constrain any further increase in cytosolic protein concentration. Consistent with this prediction, no increase in soluble protein was detected upon serum stimulation of cells at the minima of ion rhythms (36 h and 60 h, when soluble protein is already high, Fig. 2f). Serum stimulation elicited no increase in soluble protein at any timepoints in cells under mTORC inhibition. These findings indicate a bidirectional dynamic interplay between ion and protein levels: whilst changes in cytosolic protein abundance stimulate compensatory ion transport, low cellular ion content restricts the scope for acute, as well as daily, increases in protein.

### Daily regulation of SLC12A transporter activity

In concert with other transport systems, the ubiquitous and essential Na^+^/K^+^-ATPase establishes ~10-fold ion gradients over the plasma membrane, with high K^+^ and low Na^+^/Cl^−^ in the cytosol (Supplementary table 1). In response to osmotic challenge, any rapid movement of water to re-establish osmotic equilibrium and subsequent change in cell volume is rapidly countered by ion transport, which returns the cell to its original volume^17,47^. This homeostatic mechanism, known as regulatory volume increase/decrease (RVI/D), is common to all mammalian cells, and its molecular mechanisms are well characterised. K^+^ is the most abundant cytosolic osmolyte, and as such K^+^-transport *via* SLC12A cotransporters constitutes a major component of RVI/D. SLC12A family members transport Cl^−^ as obligate counterions, presumably since electrostatic constraints prevent largescale net cation efflux or import. Co-ordinated activity of SLC12A family members is thought to be regulated *via* WNK-OXSR1/SPAK1 signalling stimulated by low Cl^−^ and increased macromolecular crowding^29,48–50^ (Fig. 3a), which we confirmed using biochemical and cellular assays (Fig. 3b, c).

**Fig. 3.**
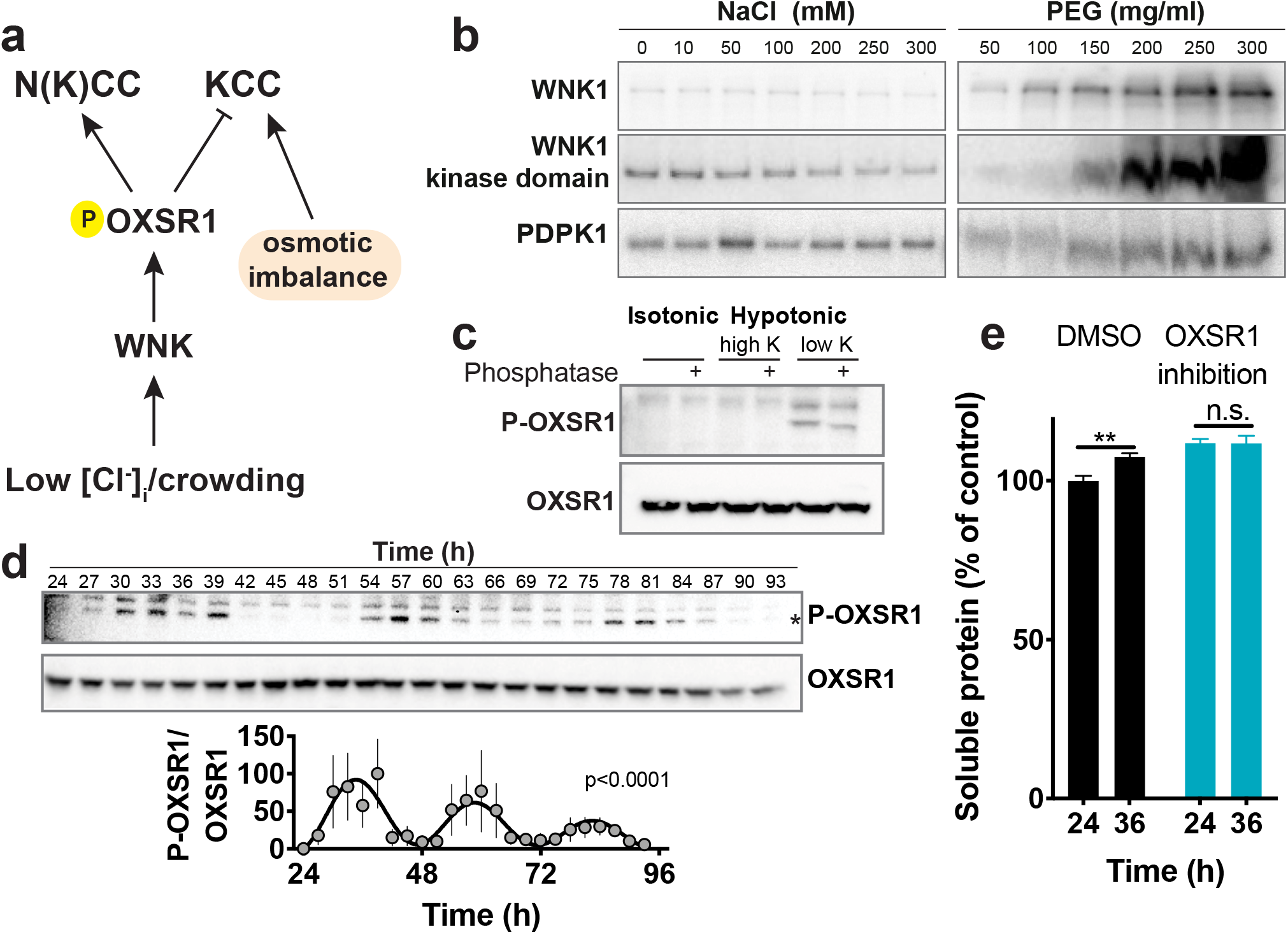
Circadian regulation of the WNK/OXSR1/SLC12A pathway activity. (a) Schematic of the WNK/OXSR1 pathway and regulation of the N(K)CC and KCC transporters. (b) Kinase activity assays for WNK1 and 3-Phosphoinositide Dependent Protein Kinase 1 (PDPK1) upon increasing concentrations of NaCl or polyethylene glycol (PEG). WNK1 but not PDPK1 is sensitive to increased macromolecular crowding (mimicked by PEG). Note that WNK1 is inhibited by high concentrations of Cl^−^. (**c**) Phosphorylation of cellular OXSR1 upon hypotonic treatment, indicating that decreased intracellular Cl^−^ increases OXSR1 phosphorylation. The addition of a phosphatase on the cell lysates confirms the identity of P-OXSR1. (**d**) Representative immunoblots and quantification showing OXSR1 and phospho-OXSR1 abundance in fibroblasts sampled every 3 h for 3 days in constant conditions (n=3). The asterisk indicates OXSR1 band. p-value indicates comparison of damped cosine wave with straight-line fit (null hypothesis = no rhythm). (**e**) Soluble protein abundance at peak and trough of protein rhythms in fibroblasts ± 30 μM of OXSR1 inhibitor closantel (n=6). Data were normalised to control at 24 h. Statistical test used was two-way ANOVA with Sidak’s MCT. Mean ± SEM shown throughout.

Low Cl^−^ and increased macromolecular crowding occur coincidentally during the circadian cycle, when mTORC1 activity is high (Fig. 2c, Supplementary Fig. 4a), leading to the prediction that WNK/OXSR1 signalling and net SLC12A-mediated ion transport should exhibit circadian regulation. Consistent with this, in cultured fibroblasts the phosphorylation profile of OXSR1 showed cell-autonomous daily rhythms (Fig. 3d), mirroring that of phospho-S6K. Peak OXSR1 phosphorylation coincided with the nadir of Cl^−^ rhythms and maximal macromolecular crowding (Supplementary Fig. 4a). Moreover, several recent phospho-proteomics studies indicate daily variation in the phosphorylation of WNK isoforms in several tissues, including cultured fibroblasts^44^, mouse liver^12^ and forebrain^27^ (Supplementary Fig. 4b-d), as well as rhythms in the phosphorylation of NKCC and KCC isoforms^27,31^, suggesting that similar mechanisms operate *in vivo*.

Our observations suggest that mTORC-dependent increases in cytosolic protein are initially facilitated by active electroneutral K^+^-Cl^−^ export via KCC, but as Cl^−^ levels fall they become rate-limiting for KCC activity. Low Cl^−^ and increased macromolecular crowding then stimulate WNK, which drives import of Na^+^ and Cl^−^ *via* OXSR1. Increased intracellular Cl^−^ availability now sustains KCC activity, whereas other transport systems export Na^+29,48,49,51^. The overall consequence of increased WNK signalling is net K^+^ export, with much smaller comparative changes in steady state Na^+^ and Cl^−^ levels. Together our data support a model whereby daily rhythms in soluble protein concentration stimulate a coordinate rhythm in SLC12A activity to maintain cytosolic osmotic potential and cell volume across the day (Supplementary Fig. 5). To test this model, we measured soluble protein abundance at the habitual peak and trough in the presence of the OXSR1 inhibitor closantel (Fig. 3e). Inhibition of OXSR1 abolished time-of-day variation in soluble protein. Interestingly, the nadir in soluble protein levels was lost in cells treated with closantel, as compared to 24 h vehicle control, consistent with SLC12A-mediated ion transport being required for return of soluble protein concentration to basal levels.

### Cell-autonomous rhythms in cardiomyocyte activity and heart rate

We next considered the physiological consequences of daily rhythms in intracellular ion content. The concentrations of Na^+^, K^+^, and Cl^−^ across the plasma membrane and their membrane permeability determine cellular resting potential and the electrophysiological properties of excitable cells^32^. We therefore employed primary mouse neonatal cardiomyocytes as a model system in which to test how cell-autonomous circadian regulation of ion abundance might impact upon cellular electrical activity. As with primary fibroblasts, we observed ~24 h cell-autonomous rhythms in intracellular Na^+^, K^+^, and Cl^−^ content (Fig. 4a, Supplementary Fig. 6a). To assess the relevance of such rhythms *in vivo,* we harvested whole heart tissue every 4 h from adult mice under diurnal conditions and analysed intracellular ion abundance by ICP-MS (Fig. 4b). We observed a significant variation in the intracellular abundance of Na^+^ and K^+^, with ion content peaking at biological times equivalent to the end of the rest phase when mTORC activity and soluble protein are normally low^10,11^.

**Fig. 4.**
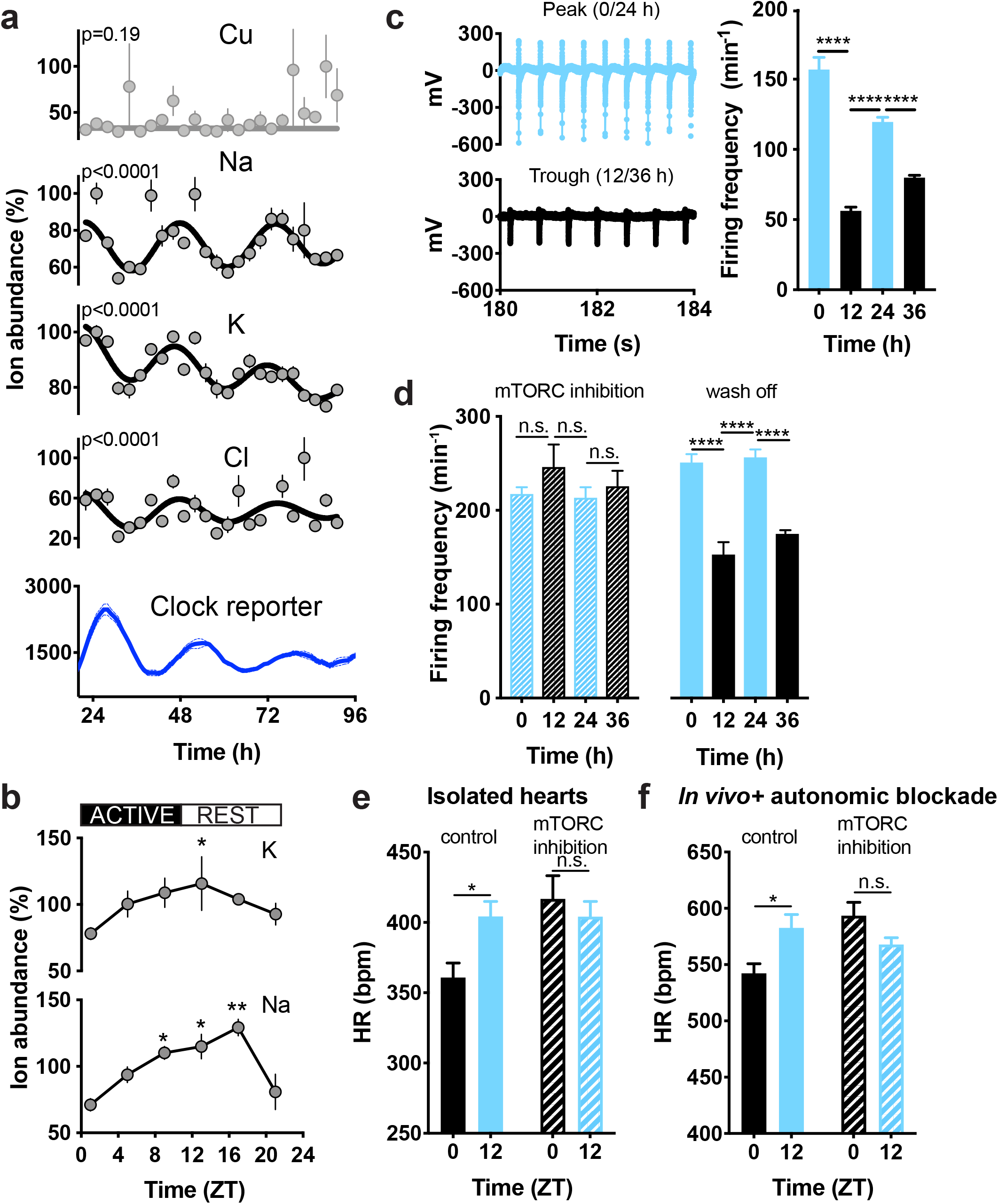
Ion fluxes facilitate circadian modulation of cardiomyocyte electrophysiology. (**a**) Cellular ion content of adult mouse heart tissue under diurnal conditions (n=3, normalised to total protein). (**b**) Abundance of selected ions and PER2::LUC reporter activity in primary cardiomyocytes (n=4) under constant conditions. Values were normalised to maximal values in each time-series. p-values indicate comparison of damped cosine wave with straight-line fit (null hypothesis = no rhythm). (**c**) Representative field potential traces of cardiomyocytes at peak or trough of ion rhythms and action potential frequency (representative biological replicate, mean signal from active electrodes is shown, n= 5, 11, 4, and 11 respectively). (**d**) Action potential frequency at peak and trough of ion rhythms in the presence of the mTOR inhibitor torin1 (50 nM), and wash off from a representative biological replicate (mean values from active electrodes are presented, n= 12, 8, 13, and 9 for torin1 and 10, 3, 3 and 5 for wash off). (**e**) Heart rate (HR) measured *ex vivo* in Langendorff-perfused hearts from control and rapamycin-treated mice collected at ZT0 and ZT12 (n=6-8/group). (**f**) HR measured *in vivo* by telemetry in control (n=6) or rapamycin-treated mice (n=5) treated with metoprolol and atropine. Time of day variation in heart rate persists under complete autonomic blockade. Mean ± SEM shown throughout. Statistical tests are one-way ANOVA with Tukey’s MCT (**b** and **c**), Two-way ANOVA with Dunnett’s MCT (**d**), one-way ANOVA and Sidak MCT (**e**), and mixed-effect analysis and Sidak MCT (**f**).

Pacemaker cardiomyocytes spontaneously depolarize and fire action potentials. Spontaneous depolarization is due to a mixed Na^+^ and K^+^ inward current (Funny-like current)^52,53^, which is affected by K^+^ and Na^+^ gradients across the plasma membrane. The slope of this slow depolarization phase, known as the pacemaker potential, determines the frequency of action potential firing. Our data predict that depolarisation should occur more quickly when intracellular ion concentrations are high, permitting firing to occur with higher frequency. To test this, we cultured primary cardiomyocytes on multielectrode arrays, and measured firing rate over two days, 12 h apart, at biological times corresponding to the peak and trough of ion rhythms. We observed a time-of-day variation in firing rate, with more frequent action potential firing when ion levels are high (Fig. 4c, Supplementary Fig. 6b). To test the relevance of reciprocal regulation of protein and ion abundance, we performed the same experiment upon inhibition of mTORC activity. We reasoned that the attenuation of protein rhythms should flatten ion rhythms and associated oscillations in cellular function. Indeed, torin1 reversibly abolished the time-of-day dependent variation in cardiomyocyte activity (Fig. 4d). Of note, the effect of torin1 on action potential firing frequency was most evident at 12 h and 36 h, when mTORC activity and soluble protein are normally high^11^. Taken together, these data are consistent with a model whereby cell-autonomous, time-of-day variation in cardiomyocyte electrical activity is driven, at least in part, by the coupling of cellular ion content to ~24 h rhythms in cytosolic protein abundance.

In line with this model, we observed time-of-day dependent variation in heart rate (HR) in Langendorff-perfused hearts from adult mice. Importantly, the time-of-day tissue-intrinsic variations in HR were abolished by mTORC1 inhibition, consistent with our findings in isolated cardiomyocytes (Fig. 4e, Supplementary Fig. 6c). Diurnal variation in HR is known to be modulated by autonomic nervous system input^54,55^, but our data in isolated cardiomyocytes and *ex vivo* hearts reveal circadian regulation by cell-autonomous mechanisms intrinsic to the heart. Concordant with this, time-of-day variation in HR was observed *in vivo* under complete autonomic blockade (*via* intraperitoneal injection of metoprolol and atropine), indicating significant modulation of HR by a circadian clock within the heart *in vivo* (Fig 4e, Supplementary Fig. 6d). More specifically, in a parallel study we found that time-of-day variations in RR and QT intervals were maintained under autonomic blockade, implying local circadian control (see also accompanying manuscript, Hayter *et al.*). Under these conditions, the daily variation in HR was also abolished by mTORC1 inhibition (Fig. 4e, f, Supplementary Fig. 6d). This result suggests that cell-autonomous mTORC1-dependent changes in cardiomyocyte ion content contribute to intrinsic HR, allowing the heart to beat faster around subjective dusk - the biological time when the greatest cardiac output is required (Supplementary Fig. 6e).

## Discussion

Maintenance of osmotic homeostasis and cell volume is a prerequisite for cell viability^19,20^, explaining the complex osmoregulatory system that protects cells from acute osmotic insults^17,28^. Very little is known about the osmotic challenge cells face over circadian time, due to intracellular variation in macromolecular concentration for example, or the mechanisms that buffer such daily oscillations and enable the cytosol to remain in osmotic equilibrium with the extracellular environment. In mouse fibroblasts, we identified cell-autonomous rhythms in soluble protein abundance and molecular crowding sustained by daily rhythms of mTORC activity, in the absence of any corresponding alteration in cell volume. By analysing the ionic composition of the cells across time, we detected oscillations in the intracellular abundance of several ions, including Na^+^, K^+^, and Cl^−^, in antiphase with soluble protein levels. Our data strongly support the hypothesis that WNK/OXSR1 signalling coordinates circadian regulation of ion influx and efflux, likely through differential regulation of SLC12A transporter activity. Whilst we do not discount that other transporters, such as LRRC8 family members^56^, may contribute to circadian ion fluxes, our observations indicate that SLC12A electroneutral ion transport acts to mitigate the osmotic challenge of daily rhythms in soluble protein abundance, thereby defending osmostasis and cell volume. Because the transport of small osmolytes buffers intracellular osmotic potential against changes in cytosolic macromolecule concentration, this confers daily variation upon the capacity of cells to respond to exogenous stimuli through changes in protein expression.

The fundamental rhythmic process we identified has important implications for the physiology of electrically active cells, such as cardiomyocytes. By performing field potential recordings at biological times corresponding to the peak and trough of ion rhythms, we indeed found a time-of-day variation in action potential firing rate of isolated cardiomyocytes. Since mTORC inhibition abolished daily firing variation in cardiomyocytes, isolated hearts, and *in vivo,* mTORC-dependent daily variation in ion content may be the major cell-autonomous factor that imparts circadian organisation to electrical activity. This highlights that soluble protein concentration can directly impact intracellular ion content, and thereby modulate cell electrophysiology.

Diurnal variation in heart rate is largely thought to be regulated by the sympathetic and parasympathetic nervous systems^57^. However, data presented here and our recent studies (Hayter *et al.*) demonstrate that time-of-day variation in heart rate occurs even under complete autonomic blockade, revealing some level of cell-autonomous regulation. Both *in vivo* and *in vitro,* the cellular increase in ion content that increases cell-intrinsic electrical activity coincides with the beginning of the active phase, when basal cardiac output must undergo the greatest shift. We suggest that cell-autonomous cardiomyocyte clocks organise the daily timing of mTORC activity and global protein synthesis/degradation so that the highest cytosolic ion levels occur in anticipation of the biological time-of-day when the heart needs to work hardest (Supplementary Fig. 6e), in concert with central autonomic control. Reduced amplitude of ion rhythms, or their temporal misalignment with sympathetic stimulation, will attenuate cardiomyocytes’ intrinsic capacity to facilitate increased heart rate at this time. We propose that dysregulation of cardiac ion or protein rhythms, during shift work or aging, renders the heart less competent to satisfy increased output demand in the early morning, contributing to the increased frequency of adverse events at this time^58–63^.

To arrive at this new understanding for the temporal regulation of cellular physiology, we focused primarily on cultured fibroblast cells, a most tractable model of the cellular mammalian clock. However, the same timekeeping mechanisms, rhythms in mTORC activity, and daily variation in ion transport have been observed in other cellular contexts, both *ex vivo* and *in vivo*^8–11,26,64^. Based on our similar recent findings in human, algal and fungal cells^9,65^, we predict that rhythms in ion transport will be observed in any eukaryotic cell with oscillations in mTORC activity, circadian or otherwise. Future work will need to explore the many potential consequences of ion transport rhythms for other aspects cellular function, as well as delineating the identity and regulation of the proteins whose cytosolic localisation changes over the course the circadian cycle.

Finally, our data posit that reciprocal regulation of ion and protein abundance is a ubiquitous cellular mechanism for osmotic homeostasis, which we expect will be of broad relevance to understanding human physiology and disease. For example, since the capacity to buffer cytosolic osmotic potential is reduced when cytosolic ion levels are low, it is tempting to speculate that this may render cells more susceptible to protein misfolding and aggregation towards the end of the daily activity cycle, when most mammals normally seek to rest and sleep.

## Materials and Methods

### Fibroblast isolation, culture, and entrainment

Lung fibroblasts were isolated from adult lung tissues^66^ of WT (C57BL6), WT PER2::LUC, and TAU PER2::LUC^43^ mice and immortalized by serial passage. Fibroblasts were cultured in DMEM high glucose (Thermo Fisher Scientific, 31966-021) supplemented with 10% serum (GE Healthcare, Hyclone™III SH30109.03) and Penicillin/Streptomycin (Pen/Strep) (Thermo Fisher), at 5% CO_2_ and 37°C.

Primary cardiac fibroblasts and cardiomyocytes were isolated from the hearts of P2-P3 neonatal mice using a commercially available dissociation kit following manufacturer’s instructions (Miltenyi Biotec, 130-098-373). In brief, heart tissue was removed and stored on ice-cold dissociation buffer (106 mM NaCl, 20 mM HEPES, 0.8 mM NaH_2_PO_4_, 5.3 mM KCl, 0.4 mM MgS0_4_, 5 mM glucose). Blood vessels and connective tissue were removed and the remaining cardiac tissue was dissected into small sections and digested. After enzymatic dissociation and enzyme inactivation, the digested tissue was filtered through a 70 μm filter (Greiner bio-one, 542070) and placed in 10 cm dishes for 2 h. The supernatant containing cardiomyocytes was collected and seeded in 96 well plates or microelectrode array chips at 1000 cells/mL density using “first day medium” (DMEM high glucose supplemented with 17% M199 (Thermo Fisher, 31150-055), 10% horse serum, 5% new born calf serum, Glutamax, Pen/Strep, and Mycozap (Lonza)). On the day after the isolation, the medium was replaced with “second day medium” (DMEM high glucose supplemented with 17% M199, 5% horse serum, 0.5% new born calf serum, Glutamax, Pen/Strep, and Mycozap). The pre-plating step allows for positive selection of primary cardiac fibroblasts that were then either kept primary (5% O_2_ incubator) or immortalized by serial passage.

Lung fibroblasts were used for Fig. 1a-d, 2c, 3d, Supplementary Fig. 1b, 2, 3c, 4b, whereas cardiac fibroblasts were used in all the remaining figures. Based on previous experiments we believe that there is no difference in circadian phenotype between fibroblasts of different tissue origin.

### Bioluminescence recordings

Cells were seeded and grown to confluence. Cells were then synchronised using temperature cycles of 12 h at 32°C followed by 12 h at 37°C for a minimum of three days. Bioluminescence was recorded using culture medium supplemented with 1 mM luciferin (and drugs where indicated). Cells were imaged using ALLIGATORs (Cairn Research) as described previously^67^, where bioluminescence was acquired for 29 min at 30-minute intervals, at constant 37°C. Mean pixel intensity form each region of interest was extracted using Fiji^68^. For period calculation, bioluminescence traces were detrended by 24 h moving average and fit using GraphPad Prism with a damped cosine wave (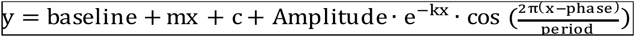) where y is the signal, x the corresponding time, amplitude is the height of the peak of the waveform above the trend line, k is the decay constant (such that 1/k is the half-life), phase is the shift relative to a cos wave and the period is the time taken for a complete cycle to occur).

### Protein extraction and quantification

For cytosolic soluble protein extraction, cells (equal number per replicate and per condition) were washed twice with phosphate-buffered saline (PBS) and incubated with a digitonin-based lysis buffer (50 mM tris pH 7.4, 0.01% digitonin, 5 mM EDTA, 150 mM NaCl, protease and phosphatase inhibitors (Roche, 4906845001 and 04693159001)) on ice for 15 mins. Only supernatant was collected (without scraping). For total protein determination, cells were washed twice with PBS and incubated with RIPA buffer (150 mM NaCl, 1% NP-40, 0.5% Na deoxycholate, 0.1% SDS, 50 mM Tris pH7.4, 5 mM EDTA, protease and phosphatase inhibitors) on ice for 15 min. After scraping, lysates were collected and sonicated using a Bioruptor sonicator (Diagenode) (30 s ON, 30 s OFF). Protein quantification was performed using intrinsic tryptophan fluorescence^69^ or BCA assay (Pierce, 23227). Reported in the figures are protein concentration values normalised as indicated in the axes or in the figure legend.

### Subcellular fractionation

Wild type PER2::LUC lung fibroblasts were seeded in 100 mm dishes (300,000 cells per dish) and grown until confluent in temperature cycles (12 h 37°C – 12 h 32°C) for 7 days. 24 h before the experiment, cells received a last medium change and were moved into constant conditions. Cytosolic, organelle and nuclear fractions were harvested every 12 h for 2 days using the Nuclei EZ Prep nuclei isolation kit (Sigma NUC-101) according to manufacturer’s instructions. Briefly, cells were washed with ice cold PBS, lysed with Nuclei lysis buffer, scraped and centrifuged at 500 g at 4°C for 5 min. Nuclei pellets were flash frozen. Supernatants were further clarified at 21,000 *g* at 4°C for 30 min to get cytosolic fractions, pellets were saved as the organelle fraction, and both fractions were flash frozen. Once thawed, nuclei and organelle pellets were resuspended in 200 μL RIPA buffer (150 mM NaCl, 5 mM EDTA pH 8.0, 50 mM Tris pH 8.0, 1% NP40, 0.5% sodium deoxycholate, 0.1% SDS, protease and phosphatase inhibitors), sonicated 4 times at high power for 30 seconds at 4°C to shear genomic DNA, and centrifuged at 21,000 *g* for 15 min at 4°C. Protein concentration was measured using BCA assay (Pierce).

### Western blotting

Equal amounts of total cell extracts were run under denaturing conditions using NuPAGE Novex 4–12% Bis-Tris gradient gels (Life Technologies). Proteins were transferred onto nitrocellulose membranes using the iBlot system (Life Technologies), with a standard (P0, 8 min) protocol. Nitrocellulose was blocked for 60 min in 5% w/w non-fat dried milk (Marvel) in Tris buffered saline/0.05% Tween-20 (TBST). Primary antibodies used were: Anti-OXSR1 (Abcam, ab125468), 1/1000; anti-Phospho-OXSR1 (Abcam, ab138655), 1/1000; anti S6K (Cell signalling, 2708), 1/2000; anti P-S6K (Cell signalling, 9205); HRP-conjugated secondary antibodies were: anti-mouse (Sigma-Aldrich, A4416) and 1/5000; anti-rabbit (Sigma-Aldrich, A6154). Chemiluminescence detection was performed using Immobilon reagent (Millipore) and the Gel-Doc™ XR^+^ system (Bio-Rad). Quantification was performed using the ImageJ gel quantification plugin or Image Lab software (Biorad). To better identify phopho-OXSR1 band, one of the replicate samples was treated with Calf intestinal alkaline phosphatase (Promega, M182A) for 60 min at 37°C.

### Mass spectrometry and Gene Ontology analysis

#### Sample preparation

Unsynchronized cells were washed twice in ice cold PBS and then lysed at room temperature with a digitonin-based lysis buffer (50 mM tris pH 7.4, 0.01% digitonin, 5 mM EDTA, 150 mM NaCl, protease and phosphatase inhibitors (Roche, 4906845001 and 04693159001)) on ice for 15 min. Supernatant was collected (without scraping) and processed for mass spectrometry analysis.

#### Enzymatic Digestion

Samples were reduced with 5 mM DTT at 56°C for 30 min and then alkylated with 10 mM iodoacetamide in the dark at room temperature for 30 min. Digestion was performed using mass spectrometry grade Lys-C (Promega) at a protein:Lys-C ratio of 100:1 (w/w) for 4 h at 37°C followed by trypsin (Promega) digestion at a ratio of 50:1 (w/w) overnight. Digestion was quenched by the addition of formic acid (FA) to a final concentration of 0.5%. Any precipitates were removed by centrifugation at 13000 rpm for 7 min. The supernatants were desalted using homemade C18 stage tips containing 3M Empore extraction disks (Sigma-Aldrich) and 2.5 mg of Poros R3 resin (Applied Biosystems). Bound peptides were eluted with 30-80% acetonitrile (MeCN) in 0.5 % FA and lyophilized.

#### Tandem mass tag (TMT) labeling

Dried peptide mixtures from each condition were re-suspended in 60 μl of 7% MeCN and 1 M triethyl ammonium bicarbonate was added to a final concentration of 200 mM. 0.8 mg of TMT10plex reagents (Thermo Fisher Scientific) was re-constituted in 41 μl anhydrous MeCN. 30 μl of TMT reagent was added to each peptide mixture and incubated for 1 hr at RT. The labeling reactions were terminated by incubation with 7.5 μl of 5% hydroxylamine for 30 min. The labeled samples were pooled into one single sample and speed Vac to remove acetonitrile. Labeled sample was desalted and then fractionated with home-made C18 stage tip using 10 mM ammonium bicarbonate and acetonitrile gradients. Eluted fractions were combined into 4 fractions, acidified, partially dried down in speed vac and ready for LC-MSMS.

#### LC MS/MS

The fractionated peptides were analysed by LC-MS/MS using a fully automated Ultimate 3000 RSLC nano System (Thermo Fisher Scientific) fitted with a 100 μm x 2 cm PepMap100 C18 nano trap column and a 75 μm×25 cm, nanoEase C18 T3 column (Waters). Samples were separated using a binary gradient consisting of buffer A (2% MeCN, 0.1% formic acid) and buffer B (80% MeCN, 0.1% formic acid), and eluted at 300 nL/min with an acetonitrile gradient. The outlet of the nano column was directly interfaced via a nanospray ion source to a Q Exactive Plus mass spectrometer (Thermo Scientific). The mass spectrometer was operated in standard data-dependent mode, performing a MS full-scan in the m/z range of 380-1600, with a resolution of 70000. This was followed by MS2 acquisitions of the 15 most intense ions with a resolution of 35000 and NCE of 33%. MS target values of 3e^6^ and MS2 target values of 1e^5^ were used. The isolation window of precursor ion was set at 0.7 Da and sequenced peptides were excluded for 40 s.

#### Spectral processing and peptide and protein identification

The acquired raw files from LC-MS/MS were processed using MaxQuant (Cox and Mann) with the integrated Andromeda search engine (v.1.6.6.0). MS/MS spectra were quantified with reporter ion MS2 from TMT 10plex experiments and searched against Mus musculus, UniProt Fasta database (March19). Carbamidomethylation of cysteines was set as fixed modification, while methionine oxidation, N-terminal acetylation and were set as variable modifications. Protein quantification requirements were set at 1 unique and razor peptide. In the identification tap, second peptides and match between runs were not selected. Other parameters in MaxQuant were set to default values.

#### Gene ontology enrichment analysis

Gene ontology enrichment analysis was performed using PANTHER 14.1 (Protein ANalysis THrough Evolutionary Relationships) Classification System ^70^. The analysis performed was a statistical enrichment test, using the Mann-Whitney U test and Benjamini-Hochberg correction for multiple comparisons.

### ^35^S incorporation assays

PER2::LUC lung fibroblasts were differentially entrained for a week with external temperature cycles in serum-free DMEM. At the end of the last cycle, cells were moved into constant conditions. A parallel plate was used for bioluminescence recording of the clock reporter (phase marker). At the indicated times, the cells were pulsed with 0.1 mCi/ml ^35^S-L-methionine/^35^S-L-cysteine mix (EasyTag™ EXPRESS35S Protein Labeling Mix, Perkin Elmer) in serum-free, Cysteine/Methionine-free DMEM for 15 min at 37°C. Afterwards, cells were washed with ice-cold PBS and lysed in digitonin-based buffer (with protease inhibitor tablet, added freshly) on ice. Equal amounts of lysates were run on 4-12% Bis-Tris SDS-PAGE using MES buffer. Gels were then dried at 80°C and exposed overnight to a phosphorimager screen. Images were acquired with Typhoon FLA700 gel scanner, and quantified using Fiji.

### Cell volume measurements

WT cardiac fibroblasts were nucleofected with pCDNA3.1 tdTomato (Neon Transfection kit, Invitrogen, MPIK10025), diluted 1 in 2 with control cells (cell nucleofected with no DNA), and seeded at high density in μslides (IBIDI, 80826). Cells were grown to confluence and entrained with temperature cycles. 24 h before the start of the recording medium was changed to “Air medium” supplemented with 1% Hyclone III, and plates sealed. Z-stacks were acquired every hour for 2.5-3 days using a Leica SP8 confocal microscope. Imaris software (Oxford Instruments) was used for 3D reconstruction, tracking and determination of cell volume. For each data set, comparison of fit was performed with GraphPad Prism. Data were fitted with either a straight line or a circadian damped cosine wave. When a circadian damped cosine wave equation was preferred, only those cells with a period between 20 - 30 h were considered rhythmic.

### Measurement of the effective diffusion rate of Quantum dots

#### Streptavidin-Quantum Dot synthesis

CdSe/CdS/ZnS core/shell Quantum Dots (QDs) were synthesized using high temperature reaction of metal carboxylates and sulfur and selenium precursors in octadecene as described previously^71^. They were transferred into water by ligand exchange with a block copolymer ligand composed of a first block of imidazole acting as anchors onto the QD surface and a second polymer block of sulfobetaine and azido-functionalized monomers^71,72^. This second block provides efficient antifouling properties from the sulfobetaine and is amenable to click chemistry mediated bioconjugation. The resulting QDs were purified using ultracentrifugation on sucrose gradients and ultrafiltration and resuspended in HBS buffer (10 mM HEPES, 150 mM NaCl pH 7.5)^72^. For streptavidin-conjugation, 10 nmol streptavidin were functionalized with dibenzylcyclooctyne (DBCO) using a 3:1 ratio of DBCO-NHS:streptavidin in borate buffer (0.1 M pH 8.0). The proteins were purified from unreacted DBCO-NHS using ultrafiltration (Vivaspin, 10 kDa cutoff, 2 rounds of 10 min, 13000 g), resuspended in HBS and mixed with 1 nmol QDs for 24 h at 4°C. Bioconjugated QDs were finally purified from unreacted proteins by ultracentrifugation in HBS (2 rounds of 25 min, 120000 g). Before delivery into cells, Streptavidin-QDs were incubated with a 15-fold molar excess of biotinylated Cell Penetrating Poly(disulfide)s (CPDs)^73^ overnight at 4°C in (0.01 M HEPES, 150 mM KCl, pH 7). On the day of the experiments QD-CPDs conjugates were diluted to a final 1 nM concentration in growth medium and added onto cells (see below). For experiment in Supplementary Fig. 2b commercial streptavidin-conjugated QDs (Thermo - Q10101MP) were used. The different size and coating of these QDs might account for the different effect size between experiments in Fig.1e and Supplementary Fig. 2b^74^.

#### Time-course experiment

WT fibroblasts were seeded in 35 mm dishes (WPI, FD3510-10), grown to confluence and differentially entrained with temperature cycles. QD motion was imaged at predicted trough and peak of protein oscillations, i.e. 24 h, 36 h, 48 h, and 60 h after medium change (DMEM 1% Hyclone III, supplemented with 10 mM HEPES to avoid any change in pH during imaging). Briefly, cells were incubated with 1 nM QD-CPD conjugates for 1 h, washed twice with PBS and imaged at 37°C, atmospheric conditions. To prevent any perturbations by media change, all treatments were performed at 37°C using the conditioned medium from the same dishes. For each time-point, we acquired several time-lapse recordings from 2-3 independent replicate dishes (between 22 and 43 movies per time point per replicate), and replicates were subsequently pooled for analysis.

#### Imaging and image processing

Imaging was performed by spinning disk confocal microscopy using a custom-built setup based on a Nikon Ti stand equipped with perfect focus system, a PLAN NA 1.45 100X objective and a spinning disk head (Yokogawa CSUX1). QDs were excited by a 488 nm laser (Coherent OBIS mounted in a Cairn laser launch) and imaged using a Chroma 595/50 emitter filter. Images were recorded with a Photometrics Prime 95B back-illuminated sCMOS camera operating in high gain mode (12 bits dynamic range), gain 1 and pseudo global shutter mode (synchronized with the spinning disk wheel). Each acquisition consisted of 1000 frames acquired at 62.5 Hz (10 ms exposure, 6 ms readout). The system was operated by Metamorph.

Estimation of the effective diffusion coefficient of QDs was performed in Fiji ^68^ and MATLAB 2017b (Mathworks) using custom codes available on request. In brief, for each field of view, QD position was precisely determined by 2D Gaussian fitting using the plugin Thunderstorm ^75^. Average localization precision across the entire dataset based on photon count was 14 ± 0.0004 nm (mean ± sem n=214, 368, 144 QDs). QD trajectories were then tracked using the MATLAB adaptation by Daniel Blair and Eric Dufresne of the IDL particle tracking code originally developed by David Grier, John Crocker, and Eric Weeks in MATLAB 2017b (http://site.physics.georgetown.edu/matlab/index.html).

For each track, the Mean Square Displacement (MSD) of segments of increasing duration (delay time 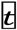) was then computed 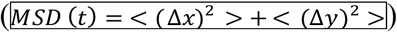 using the MATLAB class MSD Analyzer ^76^. MSD curves were then fitted to a subdiffusion model captured by the function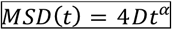^77^ with 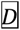 the effective diffusion rate. We then filtered the MSD curves to retain only those with a good fit (R^2^>0.9) and with 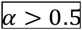 as we found this was the most accurate way to exclude immobile particles. Indeed, as shown previously ^73^, QDs delivered by our CPD technology exhibit both immobile and diffusive motion in cells, and immobile QDs have to be filtered out as they would artificially lower the estimated diffusion coefficient. We then accurately estimated the effective diffusion of these filtered tracks by fitting the first 50 points of their MSD curve by the function 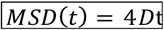. The median of the effective diffusion rate 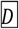 was then averaged across all the filtered tracks to give an estimate of the effective diffusion rate for a given field of view (average number of filtered tracks per field of view was always >1000). The effective diffusion rates per field of view were then average for each time point of the circadian time-course.

### Inductively coupled plasma-mass spectrometry (ICP-MS) analysis – mammalian cells

For time-course experiments, cells were seeded at high density and grown to confluence. Cells were entrained for three days with temperature cycles (12 h at 32°C, 12 h at 37°C). At the transition to the cold phase of the last cycle, medium was replaced with fresh medium and cells were moved into constant conditions (37°C). Samples were extracted every 3 h for 3 consecutive days, beginning 24 h after the medium change. Cells were washed once with ice-cold “iso-osmotic buffer 1” (300 mM sucrose, 10 mM Tris base, 1 mM EDTA, pH 7.4 adjusted with phosphoric acid (Sigma, 79614, 330-340 mOsm) and once with ice-cold “iso-osmotic buffer 2” (300 mM sucrose, 10 mM Tris base, 1 mM EDTA, pH 7.4 adjusted with acetic acid Sigma, 45727, 330-340 mOsm). Cells were then lysed with high purity 65% nitric acid (HNO_3_, Merk Millipore, 100441) supplemented with 0.1 mg/L (100 ppb) cerium (Romil, E3CE#) as a procedural internal standard. Samples were diluted in high purity water to a final concentration of 5% HNO_3_, then run using a Perkin Elmer Nexion 350D ICP-MS. Data were collected and analysed using Syngistix version 1.1. Ion concentrations were normalised for cerium abundance and analysed using GraphPad Prism. Outliers were identified and removed using the ROUT method in GraphPad Prism (Q=0.1%, high stringency). In order to determine circadian rhythmicity, data were fit by comparing a straight line with a damped cosine wave equation with the simpler model chosen unless the latter was preferred with p<0.05.

### ICP-MS analysis – heart tissue

Male C57BL/6J mice kept under 12 h/12 h light/dark (L/D) cycles were sacrificed every 4 h, at the indicated zeitgeber times (ZT). Mice were euthanized by cervical dislocation, confirmed by exsanguination. Heart tissues were removed and stored on ice in “iso-osmotic buffer 1” (see above). Tissues were diced into 3 mm slices, washed extensively with ice cold iso-osomotic buffer 1 then 2, and gently digested with an NP40-based buffer on a tube rotator (50 mM tris pH 7.5, 1% NP40, 5 mM EDTA, 50 mM choline bitartrate). Samples were spun down (21.000 g x 10 min) and the supernatant quantified using BCA (for protein quantification) and analysed by ICP-MS (for ionic composition). For each sample, ion abundance was normalised to protein abundance.

### Kinase assay

Recombinant full-length WNK1 (residues 1-2382; OriGene, RC214240) was expressed with a C-terminal Myc-DDK tag from a pCMV6-Entry backbone in Expi 293F suspension cells (Thermo Fisher Scientific, A14527). Cells were grown at 37°C, 8% CO_2_ and 125 rpm shaking in 1 L roller bottles with vented caps. A culture of 300 mL of cells at 2×10^6^ cells/mL was transfected with a mix consisting of 10 mL Expi 293 expression medium, 900 μL 1 mg/mL PEI MAX 40k (Polysciences, 24765-1) and 300 μg DNA, preincubated at room temperature for 15 min. Cells were harvested 36 hours later and spun down at 1000*g* for 5 min. Cell pellets were washed once in PBS and flash frozen. Pellets were thawed and resuspended in 20 mL lysis buffer A (50 mM Hepes pH 7.5, 125 mM KOAc, 5 mM MgAc_2_, 0.5% Triton X-100, 1 mM DTT, 1X PhosSTOP (Sigma-Aldrich, 4906845001), 1X complete EDTA-free protease inhibitor cocktail (Roche, 11873580001)), and the resulting lysate was centrifuged at 16,000 rpm for 30 min in a SS-34 rotor. The supernatant was mixed with 100 μL anti-FLAG M2 affinity resin (Millipore-Sigma, USA) and incubated at 4°C for 90 min. The beads were then washed with 9 mL of lysis buffer, 12 mL of wash buffer 1 (50 mM Hepes pH 7.5, 400 mM KOAc, 5 mM MgAc_2_, 0.1% Triton X-100, 1 mM DTT, 1X complete EDTA-free protease inhibitor cocktail), and finally 6 mL of wash buffer 2 (50 mM Hepes pH 7.5, 100 mM KOAc, 5 mM MgAc_2_, 1 mM DTT, 10% glycerol). WNK1 was then eluted from the resin in 200 μL wash buffer 2 containing 0.25 mg/mL 3X FLAG peptide.

Recombinant WNK1 kinase domain (residues 1-487) was expressed with a N-terminal His_14_-*bd*SUMO tag in *E. coli* Rosetta 2 cells (Novagen, 71397). Expression was induced with addition of 0.2 mM IPTG and cells were grown for 8 hours before harvesting. Cell pellets were resuspended via sonication in lysis buffer B (50 mM Tris pH 7.5, 300 mM NaCl, 20 mM imidazole, 5 mM DTT, 1 mM PMSF, 10% glycerol), and the lysate was then centrifuged for 45 min at 17,000 rpm in a SS-34 rotor. The resulting supernatant was mixed with Ni^2+^ affinity agarose resin (Qiagen, 30210) and incubated at 4°C for 60 min. The resin was washed with two column volumes (CV) of wash buffer 3 (50 mM Tris pH 7.5, 300 mM NaCl, 20 mM imidazole) and then with one CV of wash buffer 4 (50 mM Hepes pH 7.5, 300 mM KOAc, 2 mM MgAc_2_, 10% glycerol, 5 mM DTT). The recombinant WNK1 kinase domain was eluted via addition of 50 nM *bd*SENDP1 protease (Frey *et al*., 2014; Addgene ID 104962) in wash buffer 4.

### Recombinant PDPK1 was ordered in a purified form (abcam, ab60834)

Kinase assays were performed with 50 ng protein in assay buffer (50 mM Hepes pH 7.5, 100 mM KOAc, 2 mM MgAc_2_, 0.1 mM ATP, 0.2 mCi/mL γ-^32^P-ATP, 1X PhosSTOP) with the indicated concentrations of either NaCl or PEG 20,000 (Sigma, 81300-1KG). Reactions were incubated at 32°C for 30 min and then quenched with sample buffer and boiled. Samples were then diluted 7-fold prior to electrophoresis due to the high concentrations of PEG. Incorporation of γ-^32^P was analyzed by exposing the gel to a Phosphor screen for 24-72 hours.

### Micro electrode array (MEA) recordings

Immediately after isolation, 1×10^5^ cardiomyocytes were seeded in a small drop on the centre of the MEA chip (Multi Channel Systems, 60MEA200/30iR-Ti-gr) coated with 10 μg/mL fibronectin (Sigma, F1141). After 5 h, fresh medium was added. After 8 days in culture, with medium change performed every day, a last medium change was performed and cells kept in constant conditions. Field potential from each MEA was then recorded at predicted peaks and troughs of ion rhythms before returning cells to the incubator between time-points. Cells were maintained at 37°C during transfer to and from the incubator. Recording was done in culturing medium supplemented with 10 mM HEPES using the MEA recording device (MEA2100-2×60-System, Multi Channel Systems). All recordings were performed at atmospheric conditions with stage and custom-built heated lid held at 37°C. MEAs were allowed to equilibrate for 200 s and local field potentials from all electrodes were recorded at 10 kHz for 10 min. Data were recorded and analysed using the Multi-Channel experimenter and Multi-Channel DataManager software. Data from individual MEAs (several replicates per experiment) are shown in the main figures and in supplementary figures.

### Langendorff heart electrophysiology

Animals were culled and hearts excised between ZT0-1 or ZT11-12. Excess tissue was removed under dissection microscope in ice cold Krebs-Henseleit Buffer (KHB) solution (118 mM NaCl, 11 mM Glucose, 1.8 mM CaCl_2_, 4.7 mM KCl, 25 mM NaHCO_3_, 1.2 mM MgSO_4_, 1.2 mM KH_2_PO_4_). Hearts were cannulated through the aorta and held in place using fine thread tied around the aortic vessel. The cannula was then attached to a jacketed glass coil filled with oxygenated KHB and heart perfused at 37°C at 4 ml/min. This whole process took ~3-4 min. Monophasic action potentials were recorded using custom made silver chloride electrodes from the left atrial appendage. Any hearts that did not stabilise within the first 10 min of recording were excluded from further study. Signals were amplified and digitised using a Powerlab system (4/35) and analysed in LabChart (v8; ADinstruments, UK). After brief electrical stimulation of the right atrial appendage (600 Hz for 15s), average heart rate was calculated over 30 seconds of stable recording. Mice in the treated group were given rapamycin (dissolved in drinking water) for 2 weeks prior to culling.

### ECG telemetry implantation and analysis

Mice were implanted with telemetry devices (ETA-F20; Data Sciences International, USA) for electrocardiographic recording. Mice were anaesthetised with isoflurane, following which the telemetry remote was inserted into the abdominal cavity. Biopotential leads were secured ~1 cm right of midline at upper chest level (negative) and ~1 cm to the left of midline at the xiphoid plexus (positive). Mice were allowed a recovery period of 10 days before individual housing and the start of ECG recording. 10 second ECG waveform sweeps were collected every 5 min except for during autonomic blockade were ECG was recorded continuously. Following 5 days of ECG recording mice were given rapamycin in drinking water for 2 weeks before 5 more days of recording. At ZT0 or ZT12, mice were given 10 mg/kg metoprolol i.p. followed by 4 mg/kg atropine to achieve total autonomic blockade, confirmed by a sharp and prolonged decrease in heart rate variability (RR interval standard deviation). Intrinsic heart rate was measured over 30 min following atropine injection.

ECG analysis was carried out as previously described ^63^ using bespoke ECG software written in MATLAB (R2018a; Mathworks, USA). Briefly, for every 10s sweep R waves were detected by amplitude thresholding and ECG waveforms compared to the median of the sweep. Any that deviated significantly from this template were removed from subsequent analysis. Any sweeps with low signal/noise ratio or where >20% of detected events were excluded during template matching were excluded. Heart rate was calculated from the median RR interval of each sweep.

### Mice for Langendorff and telemetry studies

Male, C57BL/6J mice aged 10-13 weeks at the start of study, purchased from Charles River (UK), were used for ECG and Langendorff studies. Langendorff studies included some Bmal1^f/f^ animals bred in-house on the same C57BL/6J background. Mice were housed under 12:12 LD (~400/0 lux) at 22 ± 2 °C with food and water available *ad libitum*.

### Drugs

DIOA (Sigma, D129) was used at 50 μM, bumetanide (Sigma, B3023) at 100 μM, torin1 (Merck, 47599) at 50 nM, and closantel (Sigma, 34093) at 30 μM. Rapamycin (J62473, Alfa Aesar, UK) was dissolved at 50 mg/ml in ethanol, then diluted 1:1000 in water. Rapamycin solutions were kept shielded from light using tin foil and changed weekly.

### Statistical analysis

Statistical analyses were performed using GraphPad Prism. Data are presented as mean±SEM unless otherwise stated. Statistical significance was determined using two-tailed student’s t-test or ANOVA, as appropriate. Post hoc tests were used to correct for multiple comparisons as indicated in figure legend. p-values are reported using the following symbolic representation: ns = p>0.05, * = p≤0.05, ** = p≤0.01, *** = p≤0.001, **** = p≤0.0001.

All animal experiments were licensed under 1986 Home Office Animal Procedures Act (UK) and carried out in accordance with local animal welfare committee guidelines.

## Supplementary Materials

**Supplementary Fig. 1**. Gene ontology analysis of digitonin lysates and validation of QDs.

**Supplementary Fig. 2**. mTORC inhibition attenuates soluble protein rhythms and time of day variation in the diffusion coefficient of Quantum Dots.

**Supplementary Fig. 3**. Ion rhythms in fibroblasts occur with no volume change.

**Supplementary Fig. 4** Rhythms in the WNK/OXSR1 pathway.

**Supplementary Fig. 5**. Working model.

**Supplementary Fig. 6**. Ion rhythms in cardiomyocytes and firing frequency.

**Supplementary Fig. 7**. Western blots related to Supplementary Fig. 2a.

**Supplementary Fig. 8**. Western blots related to Fig. 3.

**Supplementary table 1**. Intracellular and extracellular ion concentrations in mammalian cells.

**Movie S1**. Tracking and effective diffusion of quantum dots at peak and trough of protein rhythms.

**Movie S2**. Tracking and effective diffusion of quantum dots upon challenge with media at different osmolality.

**Hayter *et al.***

## Acknowledgments

We thank Alex Harmer, Helen Causton, and past and present O’Neill lab members for valuable discussion and contribution, particularly Priya Crosby and Ned Hoyle, visual aids, and the biological services group for assistance with animal work and husbandry.

## Funding

E.D. was supported by the Medical Research Council (MC_UP_1201/13) and the Human Frontier Science Program (CDA00034/2017-C); RSE by a Wellcome Trust Sir Henry Dale Fellowship (208790/Z/17/Z). This work was supported by the AstraZeneca Blue Skies Initiative and the Medical Research Council (MC_UP_1201/4).

## Author contributions

J.S.O., P.N., R.S.E. and A.S. conceived the idea and wrote the manuscript. E.B and S.M. synthesised the cell penetrating poly(disulphide)s, T.P. and N.P. synthesised quantum dots. A.S., D.W., S.B., J.W., A.Z., E.S., S.P.C., A.B., E.H., A.G, A. I., J.D., R.V., D.B., E.D. and R.S.E. performed experiments and analysis, G.v.O. made pivotal intellectual contributions.

## Competing interests

P.N. holds shares in AstraZeneca. The other authors declare no competing interests.

## Data and materials availability

All data is available in the main text or the supplementary materials. Codes for quantum dot tracking and analysis will be available on request.

## Supplementary Figures

**Supplementary Fig. 1.**
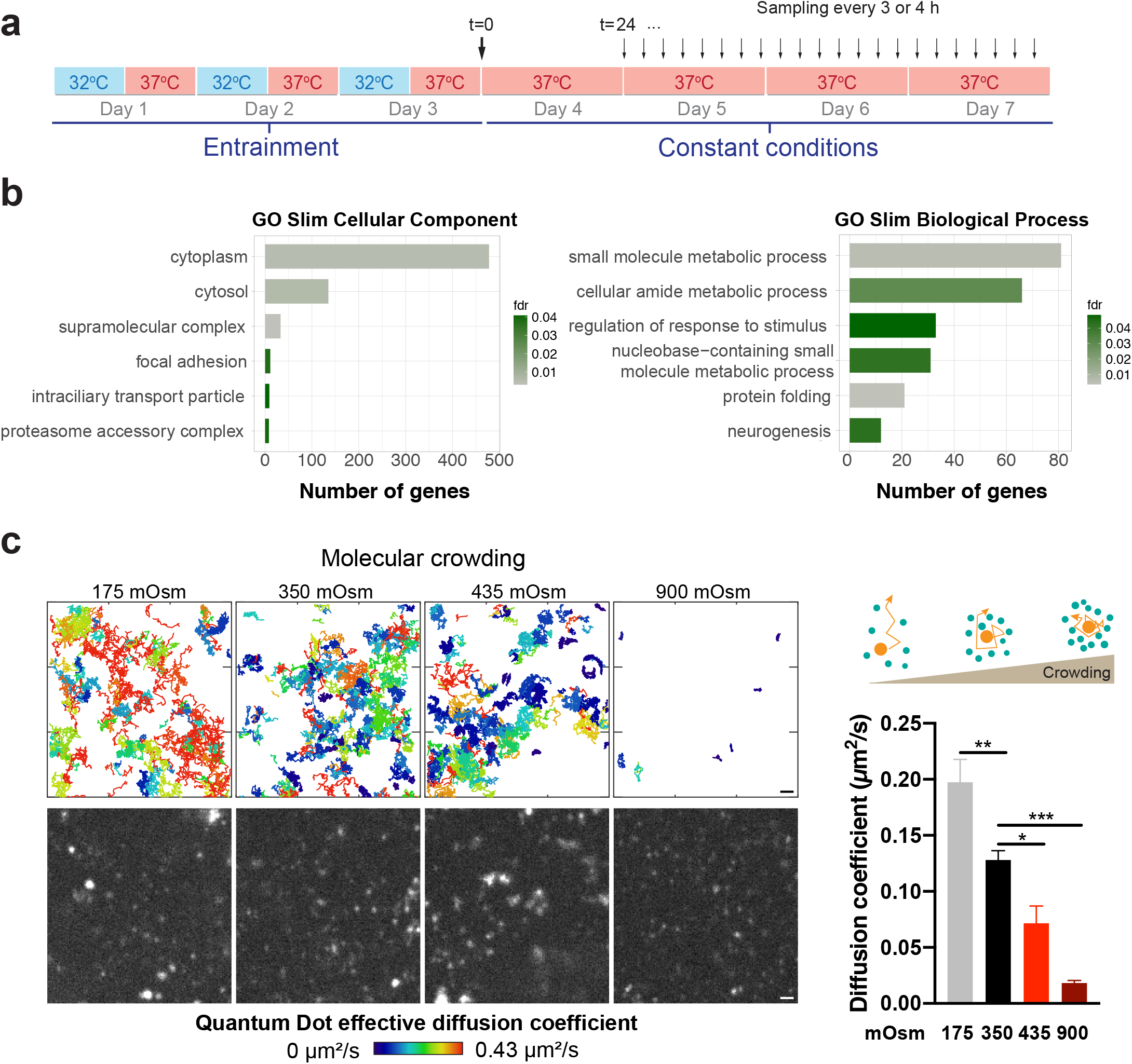
Gene ontology analysis of digitonin lysates and validation of QDs. (**a**) Schematics of time course experiments. (**b**) Gene ontology analysis on digitonin extracts (n=4) using PANTHER enrichment test. Shown are top 6 terms for GO-Slim Cellular Component and GO-Slim Biological Process annotation data categories. Mann-Whitney U test and Benjamini-Hochberg correction (FDR, p<0.05) were performed. (**c**) Validation of diffusion quantification from the effective diffusion rate of quantum dots (QDs) in fibroblasts upon treatment with media of different osmolality (n=10, 16, 5, 5) and representative tracking images. Data are presented as Mean ± SEM. Statistical test used was one-way ANOVA with Dunnet’s MCT.

**Supplementary Fig. 2.**
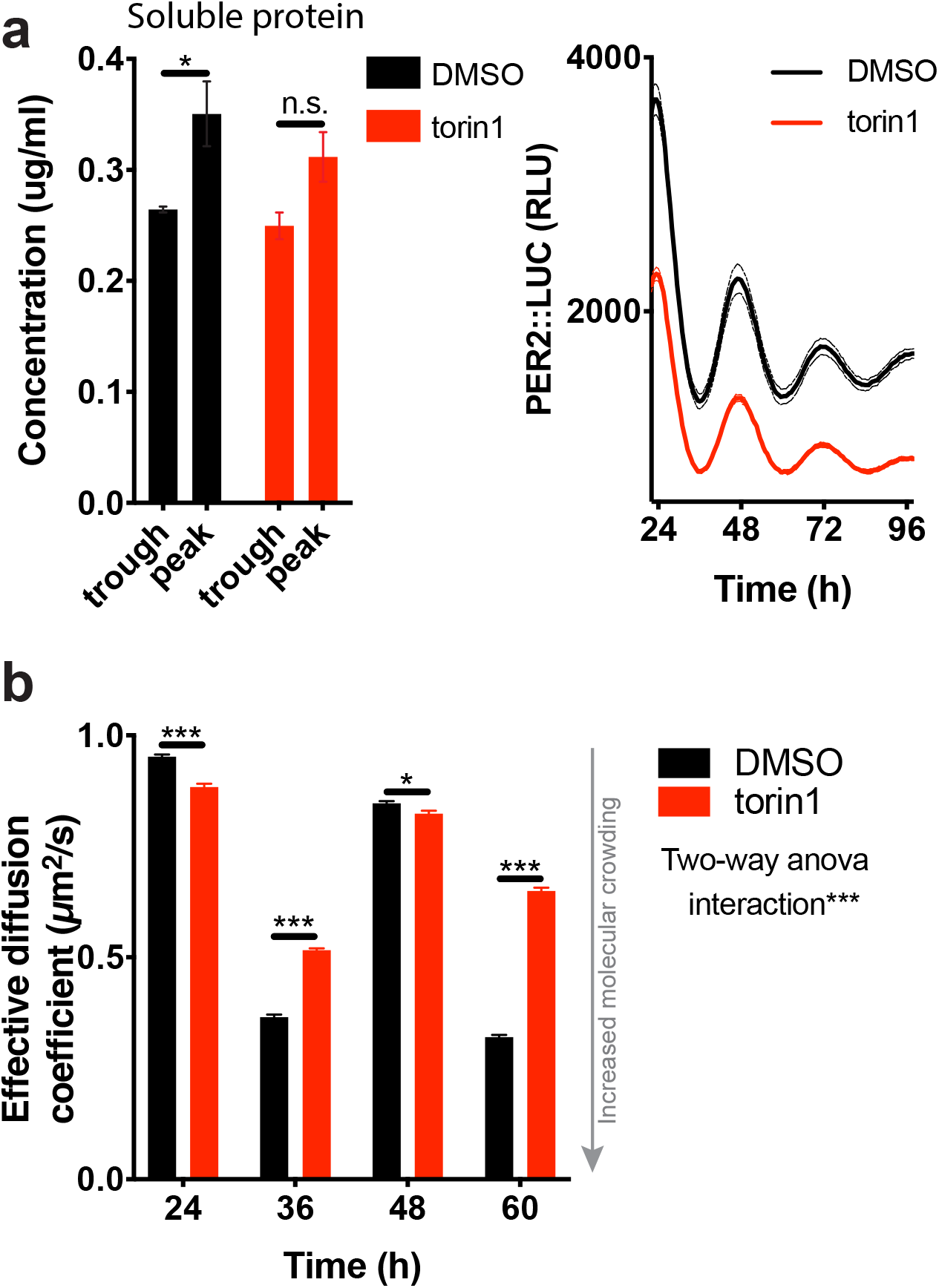
mTORC inhibition attenuates soluble protein rhythms and time of day variation in the diffusion coefficient of Quantum Dots. (**a**) Quantification of soluble protein (n=3) in fibroblasts ± 50 nM torin1 at peak and trough of protein rhythms and bioluminescence data of the circadian clock reporter (n=3). (**b**) Effective diffusion rate of QDs in fibroblasts ± 50 nM torin1. Data are presented as mean ± SEM. Statistical significance calculated using one-way ANOVA with Sidak’s MCT in (**a**) and two-way ANOVA in (**b**).

**Supplementary Fig. 3.**
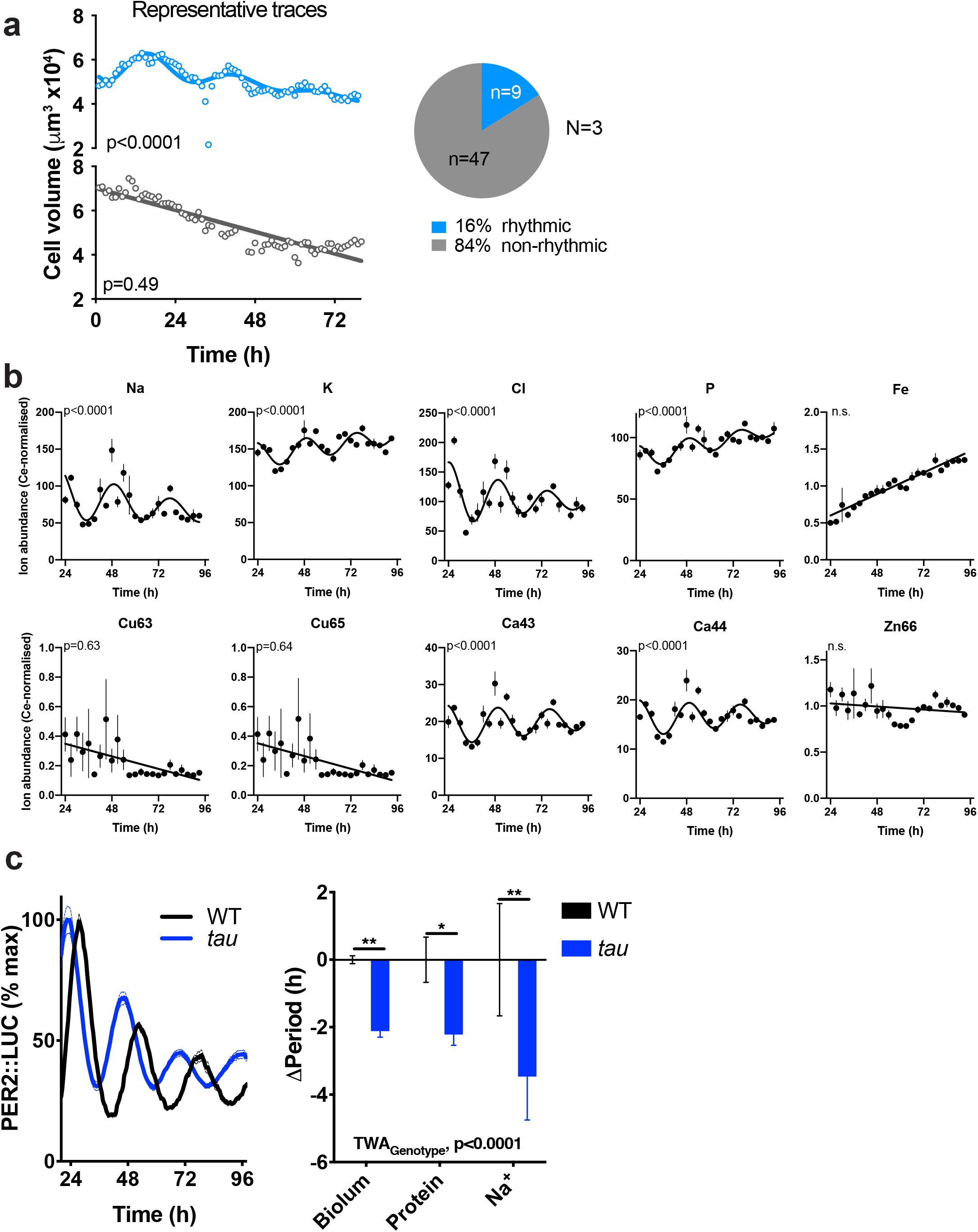
Ion rhythms in fibroblasts occur with no volume change. (**a**) Representative measurements and quantification of fibroblast volume over several circadian cycles reveals that most (84%) cells show no significant circadian variation in volume. While feeding-dependent diurnal variation in hepatocyte volume was recently reported ^6^, only a small proportion (16%) of individual fibroblasts showed significant ~24 h variation in volume i.e. where a damped sine wave fit was preferred (p<0.05) over a straight-line fit (null hypothesis = no rhythm). Thus, changes in cellular volume cannot account for the daily variations in ion and cytosolic protein abundances we observe. (**b**) ICP-MS quantification of biologically-relevant ions in primary fibroblasts (n=4). p-values indicate comparison between a damped cosine wave fit compared with a straight-line (null hypothesis, no rhythm). (**c**) Bioluminescence control for WT and tau mutant fibroblasts. Comparison of ΔPeriod for clock reporter, soluble protein rhythms, and Na^+^ rhythms. Statistical test used was 2-way ANOVA and Holm-Sidak’s MCT. Data are presented as mean ± SEM.

**Supplementary Fig. 4.**
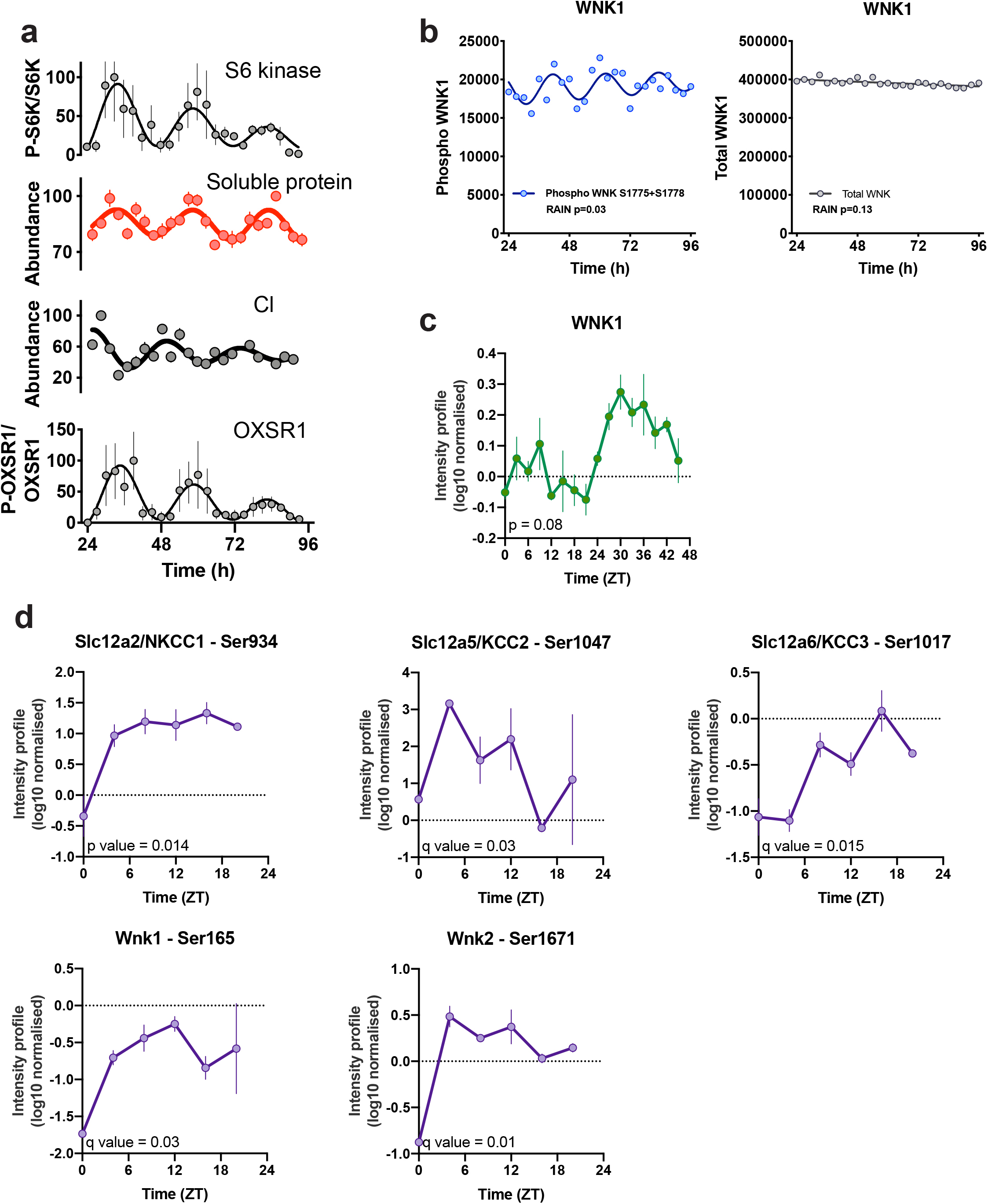
Rhythms in the WNK/OXSR1 pathway. (**a**) Composite figure summarizing S6 Kinase, soluble protein, Cl, and OXSR1 rhythms, and phase relationship. Intensity profiles (phosphorylation) of indicated proteins taken from Wong *et al.*^44^ (**b**), Robles *et al.*, 2017^12^ (**c**), and Brüning *et al.*, 2019^27^ (**d**).

**Supplementary Fig. 5.**
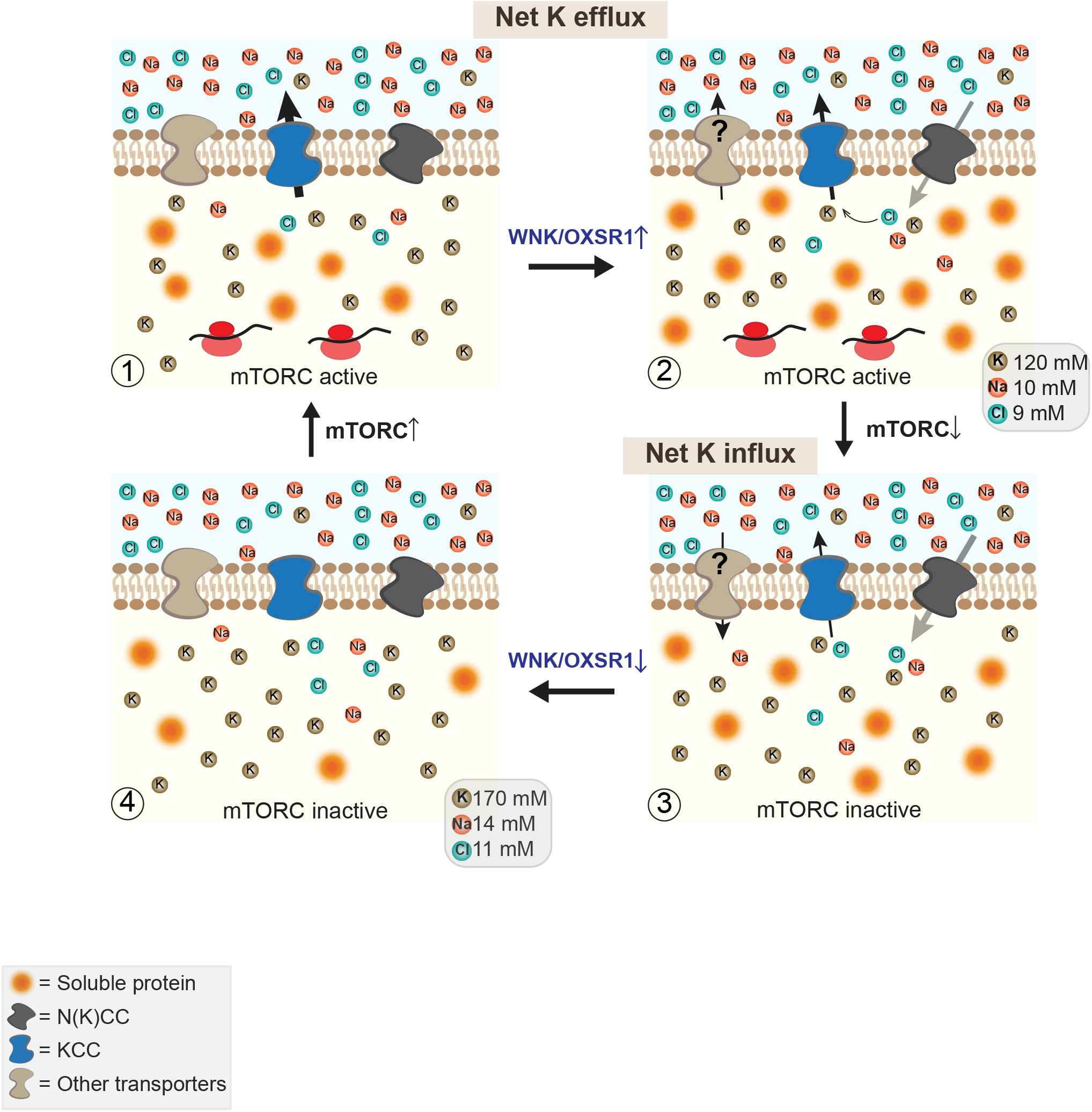
Working model. Schematic of working model, from top left: (1) mTORC activation facilitates increased macromolecular crowding in the cytosol, whose osmotic potential is buffered by electroneutral KCl export; (2) increased crowding and reduced intracellular Cl^−^ concentration stimulate increased N(K)CC activity *via* WNK/OSXR1 to facilitate further net K^+^ efflux through Cl^−^ recycling. (3) mTORC inactivation elicits net K^+^ influx to buffer the fall in cytosolic protein until (4) a new steady state is achieved and N(K)CC inactivated. Whilst SLC12A family members clearly contribute to this osmotic buffering system, other transporters are likely involved.

**Supplementary Fig. 6.**
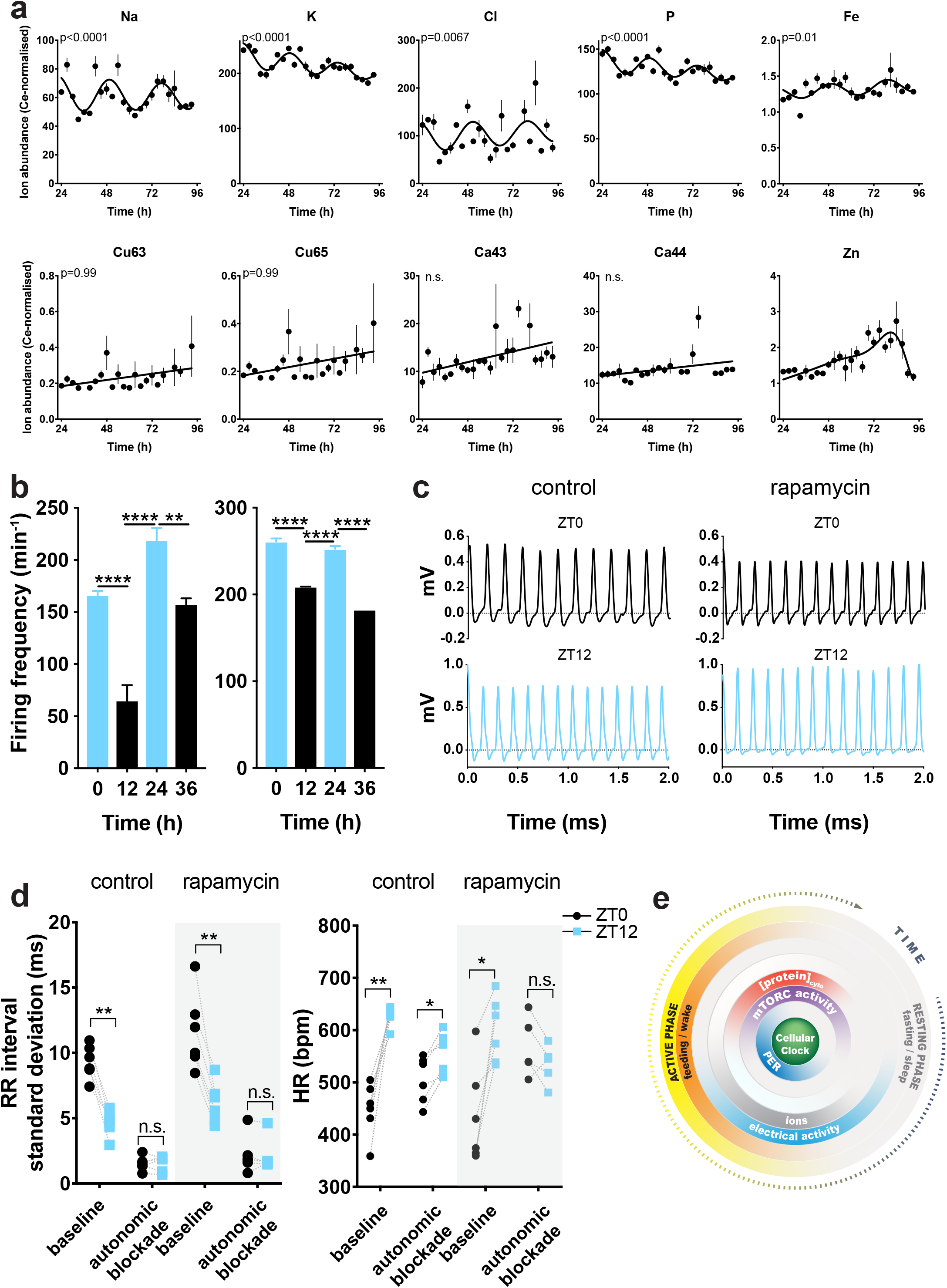
Ion rhythms in cardiomyocytes and firing frequency. (**a**) ICP-MS quantification of biologically-relevant ions in primary cardiomyocytes (n=4). p-values indicate comparison between a damped cosine wave fit compared with a straight-line (null hypothesis = no rhythm). (**b**) Firing frequency of primary cardiomyocytes at the peak and trough of ion rhythms in two more independent biological replicates (mean values from active electrodes are presented). Statistical significance was calculated using one-way ANOVA with Tukey’s test. (**c**) Representative traces of *ex vivo* Langendorff recordings (monophasic action potentials) from control or rapamycin-treated mice at indicated times. (**d**) RR interval standard deviation and Heart rate (HR) before and after autonomic blockade in control and rapamycin-treated mice. Reduction in RR interval standard deviation confirms the effect of autonomic blockade. Note that this does not affect the time-of-day variation in HR in control mice but only in the rapamycin-treated group. (**e**) Summary cartoon of the temporal relationships for *ex vivo* measurements with respect to gene expression and behaviour *in vivo*. Data are presented as mean ± SEM.

**Supplementary Fig. 7.**
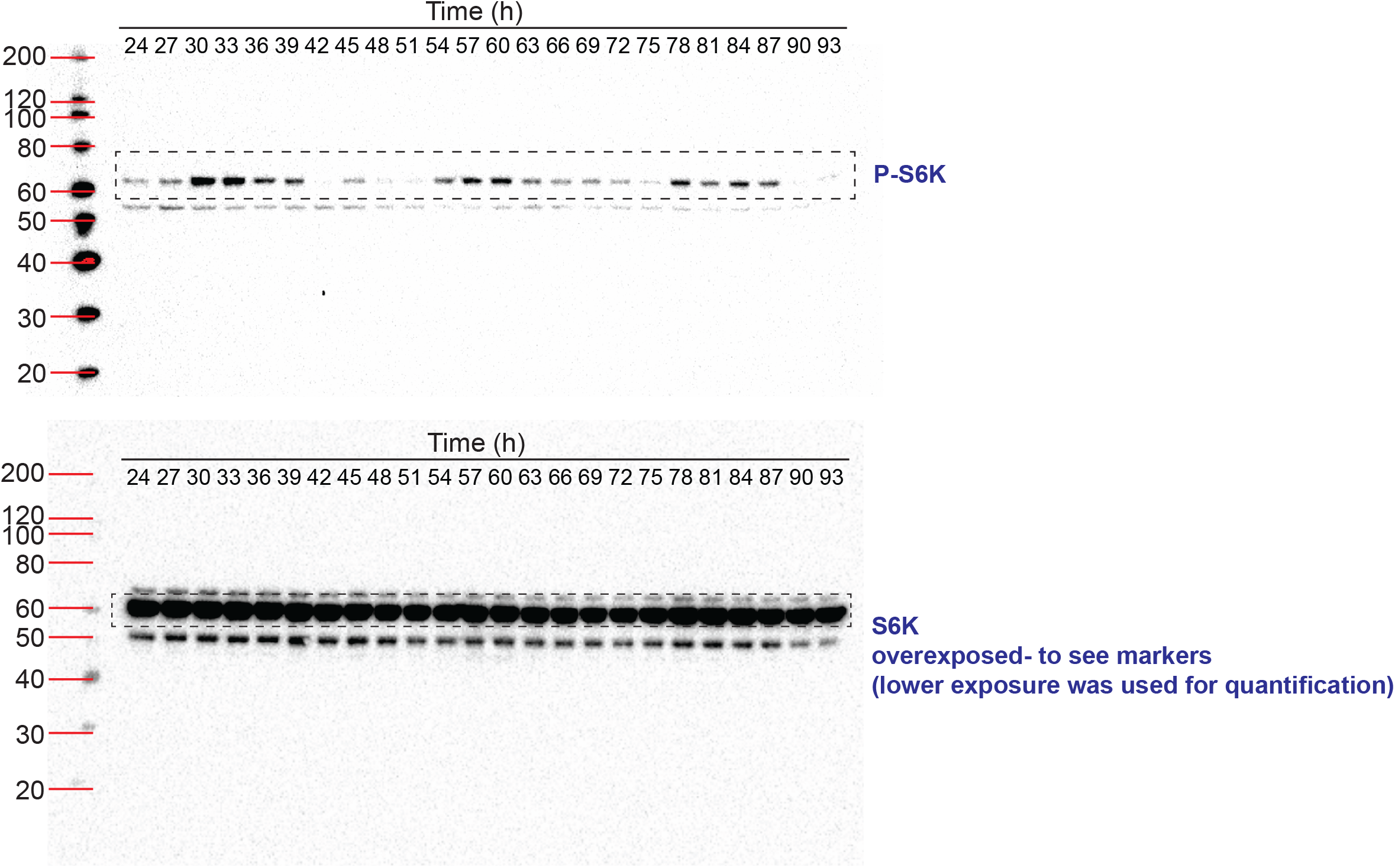
Western blots related to Fig. S2A. Uncropped original immunoblots of SDS-page gels presented in Fig. 1a.

**Supplementary Fig. 8.**
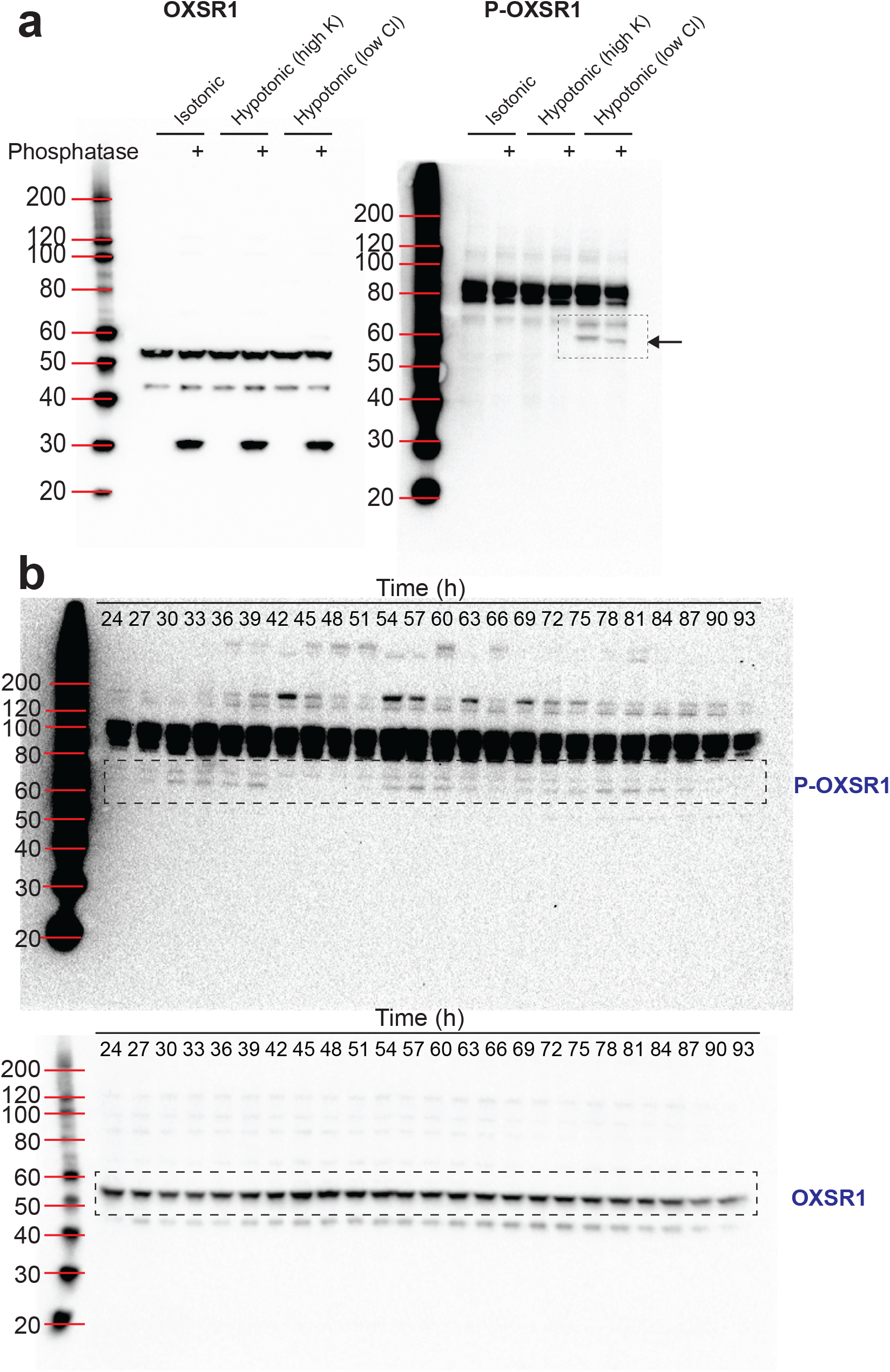
Western blots related to Fig. 3. (**a**) Uncropped original immunoblots of SDS-page gels presented in Fig. 3c showing OXSR1 And Phospho-OXSR1. Phosphorylation of OXSR1 (just above 60 Kb) is increased upon treatment with a hypotonic low chloride solution (67.5 mM sodium gluconate, 2.5 mM potassium gluconate, 0.25 mM CaCl_2_, 0.25 mM MgCl_2_, 0.5 mM Na_2_HPO_4_, 0.5 mM Na_2_SO_4_ and 7.5 mM Hepes (pH 7.5). Treatment of the lysate with alkaline phosphatase induces a shift in the molecular weight of the band. (**b**) Uncropped original immunoblots of SDS-page gel presented in Fig. 3d.

## Supplementary Materials

**Table 1.**
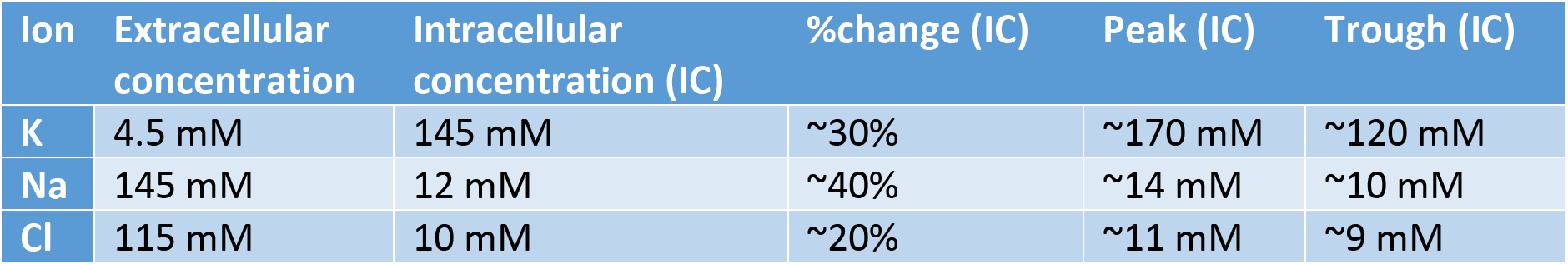
Intracellular and extracellular ion concentrations in mammalian cells. Intracellular and extracellular ion concentrations in mammalian cells from ^16^. Calculation of predicted intracellular concentration at peak and trough of ion rhythms using the Goldman– Hodgkin–Katz equation (% change calculated from data in Fig. 2b).

| Ion | Extracellular concentration | Intracellular concentration (IC) | %change (IC) | Peak (IC) | Trough (IC) |
| --- | --- | --- | --- | --- | --- |
| K | 4.5 mM | 145 mM | ~30% | ~170 mM | ~120 mM |
| Na | 145 mM | 12 mM | ~40% | ~14 mM | ~10 mM |
| Cl | 115 mM | 10 mM | ~20% | ~11 mM | ~9 mM |

**Supplementary Video 1, Related to Figure 1e**.

Tracking and effective diffusion of quantum dots at peak and trough of protein rhythms.

**Supplementary Video 2, Related to Supplementary Fig. 1c**.

Tracking and effective diffusion of quantum dots upon challenge with media at different osmolality.

## Notes

### Competing Interest Statement

Peter Newham holds shares in AstraZeneca.

